# The genetic composition of hybrid *Mangifera*

**DOI:** 10.1101/2023.03.27.533847

**Authors:** Emily J. Warschefsky, Eric J.B. von Wettberg

## Abstract

Homoploid hybridization is known to play an important role in the evolution of plants, including many crop species, but can have different outcomes including introgression between parental taxa and the formation of new evolutionary lineages. We investigate the occurrence and consequences of hybridization between the economically important tree crop *Mangifera indica* (mango) and two congeneric species in Southeast Asia. A total of 90 samples of the hybrid *M. odorata* and its parental taxa, *M. indica* (mango) and *M. foetida*, along with 65 samples of a newly proposed hybrid, *M. casturi* and its putative parental taxa, *M. indica* and *M. quadrifida*, were sampled and genotyped using restriction site associated DNA sequencing. For each hybrid, we assessed population structure and admixture and indices of genetic diversity, including multilocus linkage disequilibrium. We found no evidence of introgression between *M. foetida* and *M. indica* cultivars from Southeast Asia, but find support for a hybrid origin of *M. casturi*. Both hybrids show low levels of intraspecific genetic diversity and individuals have high genetic identity and significant multilocus linkage disequilibrium. For both *M. odorata* and *M. casturi*, our results are consistent with hybrid lineages that have formed only a few times and have since been maintained clonally. While grafting may play a role in the continued propagation of these hybrids, we suggest that the ability of *M. odorata* and *M. casturi* to reproduce asexually through nucellar polyembryony has allowed the hybrids to persist independently of grafting.

## Introduction

Hybridization has long been considered a creative force in the evolutionary process and is known to have played a particularly important role in plant evolution and speciation (e.g., Stebbins, 1950; Anderson and Stebbins, 1954; Arnold, 1992; Rieseberg, 1997; Mallet, 2007; Soltis and Soltis, 2009; Abbott et al., 2013; Yakimowski and Rieseberg, 2014). Broadly defined as the interbreeding of individuals from different genetic populations, hybridization occurs via multiple mechanisms and results in varied outcomes (e.g., Barton and G.M., 1985; Mallet, 2007; Soltis and Soltis, 2009; Abbott et al., 2013). In plants, interspecific hybridization can occur with or without whole genome duplication, resulting in homoploid or allopolyploid hybrids, respectively. Typically, allopolyploids are reproductively isolated from their parental taxa as a result of chromosomal incompatibilities, and are therefore generally considered cases of saltational evolution (Mallet, 2007; Soltis and Soltis, 2009). However, the consequences of homoploid hybridization are more complex. In some cases, low fitness of homoploid hybrids is thought to contribute to reproductive isolation of parental taxa (Servedio and Noor, 2003), while in other cases, hybrid intermediates may provide a bridge for introgression (gene flow) between parental taxa (e.g., Anderson, 1949; Arnold, 1992; Arnold et al., 2012). Alternatively, homoploid hybrids may form lineages (or species) that are ecologically distinct and which are or may become reproductively isolated from parental species (Soltis and Soltis, 2009; Yakimowski and Rieseberg, 2014).

Although hybridization is a widespread phenomenon in plants, it is known to be more common in certain circumstances. For instance, long-lived plants, which tend to be self-incompatible, are more likely to hybridize than annuals (Ellstrand et al., 1996; Petit and Hampe, 2006; Miller and Gross, 2011). Closely related allopatric species that are brought into secondary contact either by natural dissolution of geographic barriers or by human-mediated introductions are also more likely to hybridize, as they may not have developed reproductive isolation (Coyne and Orr, 2004; Harrison and Larson, 2014). Therefore, it comes as no surprise that many domesticated plants, especially perennials, are of confirmed or putative hybrid ancestry (Warschefsky et al., 2014), including apple (*Malus* x*domestica*, Coart et al., 2006; Cornille et al., 2012), potato (*Solanum tuberosum*, Rodríguez et al., 2010), banana (*Musa acuminata*, De Langhe et al., 2010), and many citrus species (*Citrus* spp. Wu et al., 2018). Such hybrid crops may be the inadvertent result of cultivation in proximity to congeneric species or the outcome of intentional crosses.

The genus *Mangifera* includes approximately 69 tropical tree species, all of which are native to South and Southeast Asia (Kostermans and Bompard, 1993). The most well known species of *Mangifera, M. indica* (mango), was likely domesticated around 4,000 years ago in India (Mukherjee, 1949). The first introduction of *M. indica* outside its original center of domestication likely occurred during the 4th and 5th centuries, when Buddhist monks from India traveled to Southeast Asia (Mukherjee, 1949). There, it came into contact with approximately 35 other species of *Mangifera* native to the region, many of which are edible and a few of which are regionally cultivated in small orchards and backyard gardens (Kostermans and Bompard, 1993).

Though only *M. indica* has been investigated, *Mangifera* species are assumed to be self-incompatible and therefore obligately outcrossing (Kostermans and Bompard, 1993; Mukherjee and Litz, 2009). It follows that the cultivation of such outcrossing congers in close proximity to one another would lead to interspecific hybridization, a phenomenon that has been documented in the genus (Kostermans and Bompard, 1993; Teo et al., 2002; Bompard, 2009).

Today, *M. indica* is cultivated in subtropical and tropical regions around the world and is one of the most economically important tropical fruit species (FAO, 2003; FAOSTAT, 2013). On the basis of fruit morphology, cultivars of *M. indica* are separated into two types, Indian and Southeast Asian (Crane and Campbell 1994). Molecular research demonstrates that these cultivar types form distinct genetic groups (Schnell et al., 2005; Dillon et al., 2013), and that Southeast Asian cultivars contain genetic diversity that is not present in Indian populations (Warschefsky and von Wettberg 2019). However, potential causes of the genetic and morphological differentiation observed in *M. indica* have not been explored. While one theory is that Indian and Southeast Asian cultivar types are the result of independent domestication events (Bompard, 2009), an alternative explanation is that the novel diversity observed in Southeast Asian *M. indica* was introduced by introgressive hybridization with a congeneric species.

The kuwini mango, *M. odorata*, was first described by Griffith in 1854 (Griffith, 1854). Only known from cultivation in Southeast Asia (Kostermans and Bompard, 1993; Teo et al., 2002), *M. odorata* had long been thought to be a hybrid of *M. indica* and *M. foetida* (horse mango), another cultivated species, on the basis of morphological characters (Hou, 1978). In 2002, Teo et al. confirmed the hybrid status of *M. odorata* using amplified fragment length polymorphisms (AFLPs) and found that *M. odorata* has greater genetic affinity to *M. foetida* than to *M. indica*. The species was later given taxonomic status as a hybrid, *M.* x*odorata* Griff, (pro sp.) (Kiew, 2002). However, no further research has explored the dynamics of hybridization between *M. indica* and *M. odorata* or tested whether the novel diversity observed in Southeast Asian cultivars of *M. indica* is the result of introgression from *M. foetida*.

*Mangifera casturi*, (kastooree mango) was recently described by Kostermans and Bompard (1993). Only known from cultivation in Kalimantan, the Indonesian region of Borneo, *M. casturi* is classified as extinct in the wild (IUCN, 2012). The species’ characteristic dark purple to black fruit and orange flesh led the authors to suggest it was most closely related to another Bornean species with similar fruit coloration, *M. quadrifida.* However, recent phylogenetic analysis indicates that despite its fruit morphology, *M. casturi* is not closely allied to *M. quadrifida*, but is instead a close relative of *M. indica* (Warschefsky 2018). Considering this new phylogenetic information, it is feasible that *M. casturi* may be a hybrid of *M. indica* and *M. quadrifida*, an idea not previously proposed.

Here, we explore the occurrence and consequences of hybridization in two separate instances in *Mangifera*. Analyzing SNP markers obtained by RAD sequencing, we: 1) determine whether the novel genetic diversity observed in Southeast Asian *M. indica* cultivars can be attributed to introgression from *M. foetida* via *M. odorata*; 2) look for evidence of a hybrid origin of *M. casturi*; and 3) characterize the genetic diversity and population structure of *M. odorata* and *M. casturi*.

## Methods

### Sampling

Leaf samples were collected from 280 specimens in living collections in Singapore, Indonesia, Malaysia, and the United States. Fresh leaves were stored at -80°C or dried in silica and stored at 4°C. Genomic DNA was extracted using a Qiagen DNEasy mini extraction kit or by a modified CTAB protocol (Doyle and Doyle, 1990). Samples were analyzed as two independent datasets: 90 samples were analyzed to examine hybridization in *M. odorata* (Table 1), and 65 individuals were analyzed to examine hybridization in *M. casturi* (Table 2).

**Table 1.**
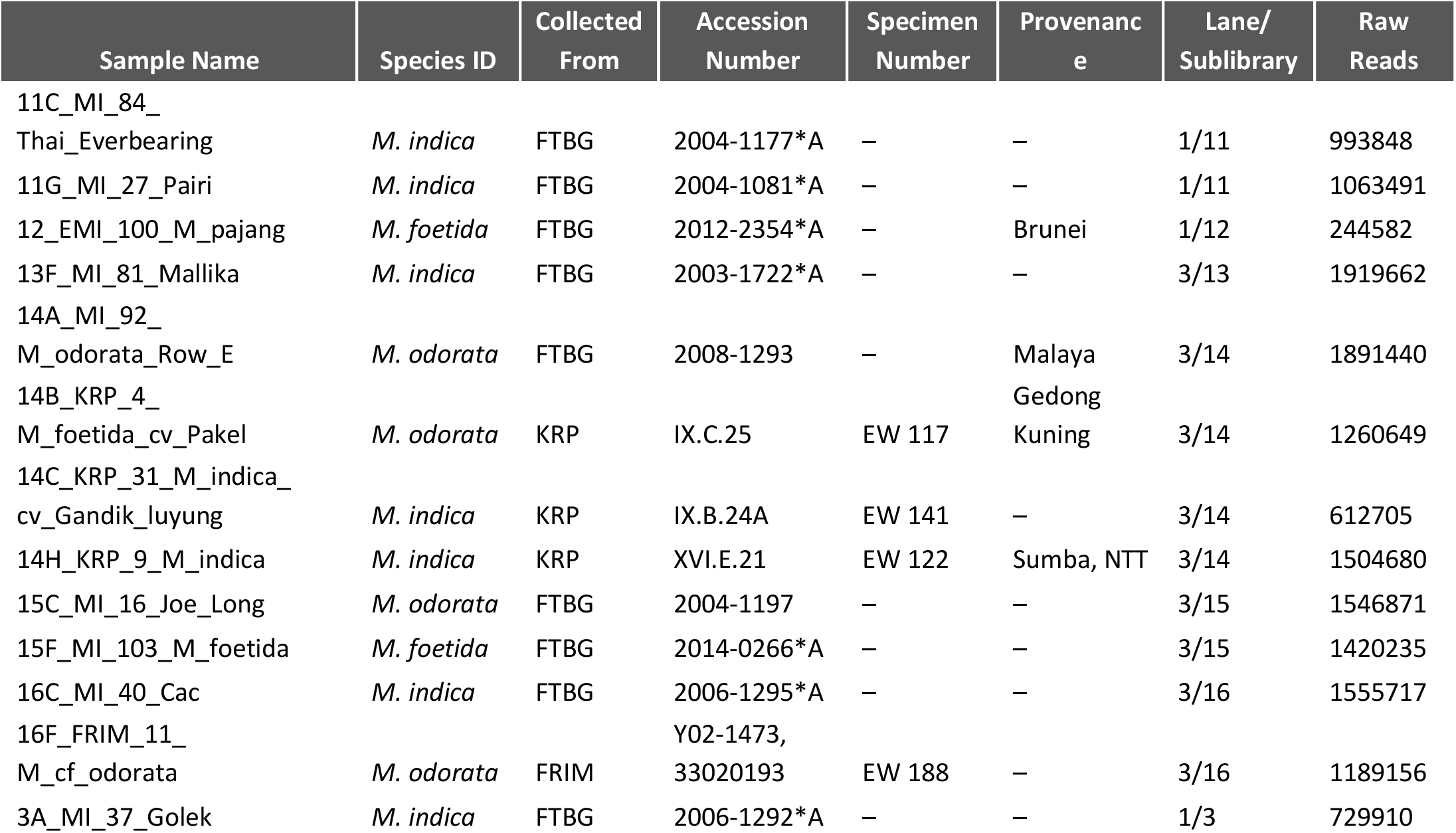
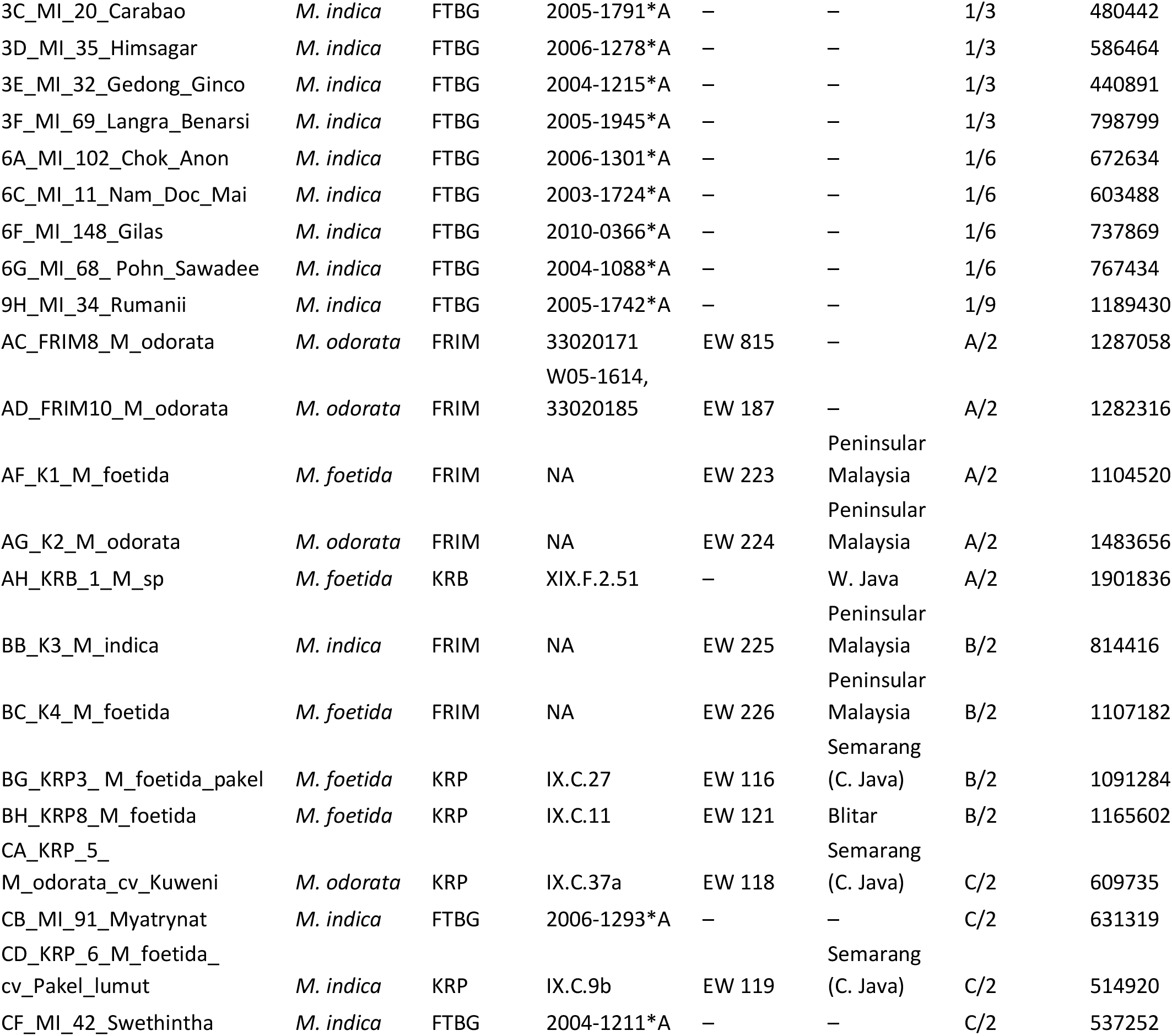
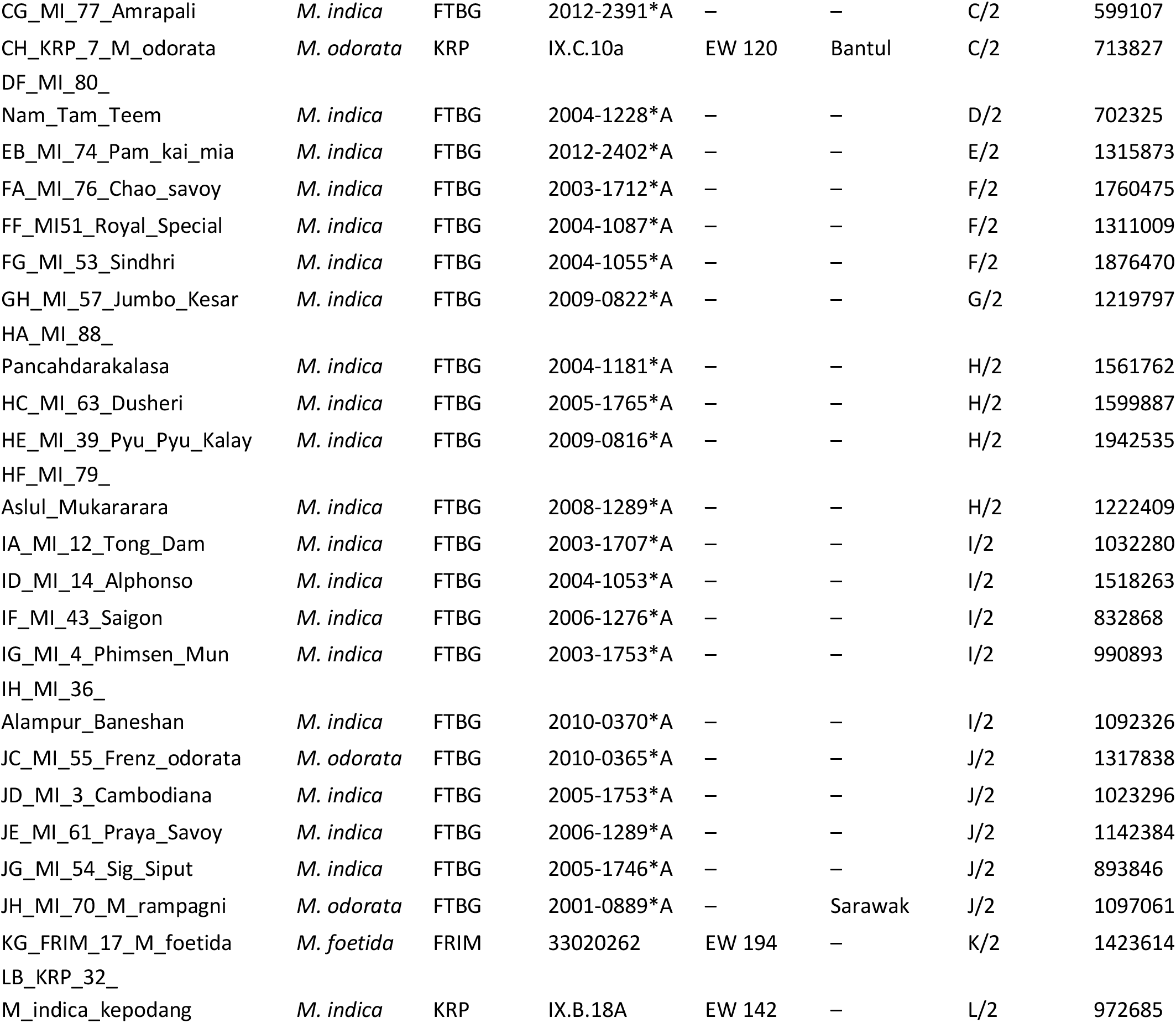
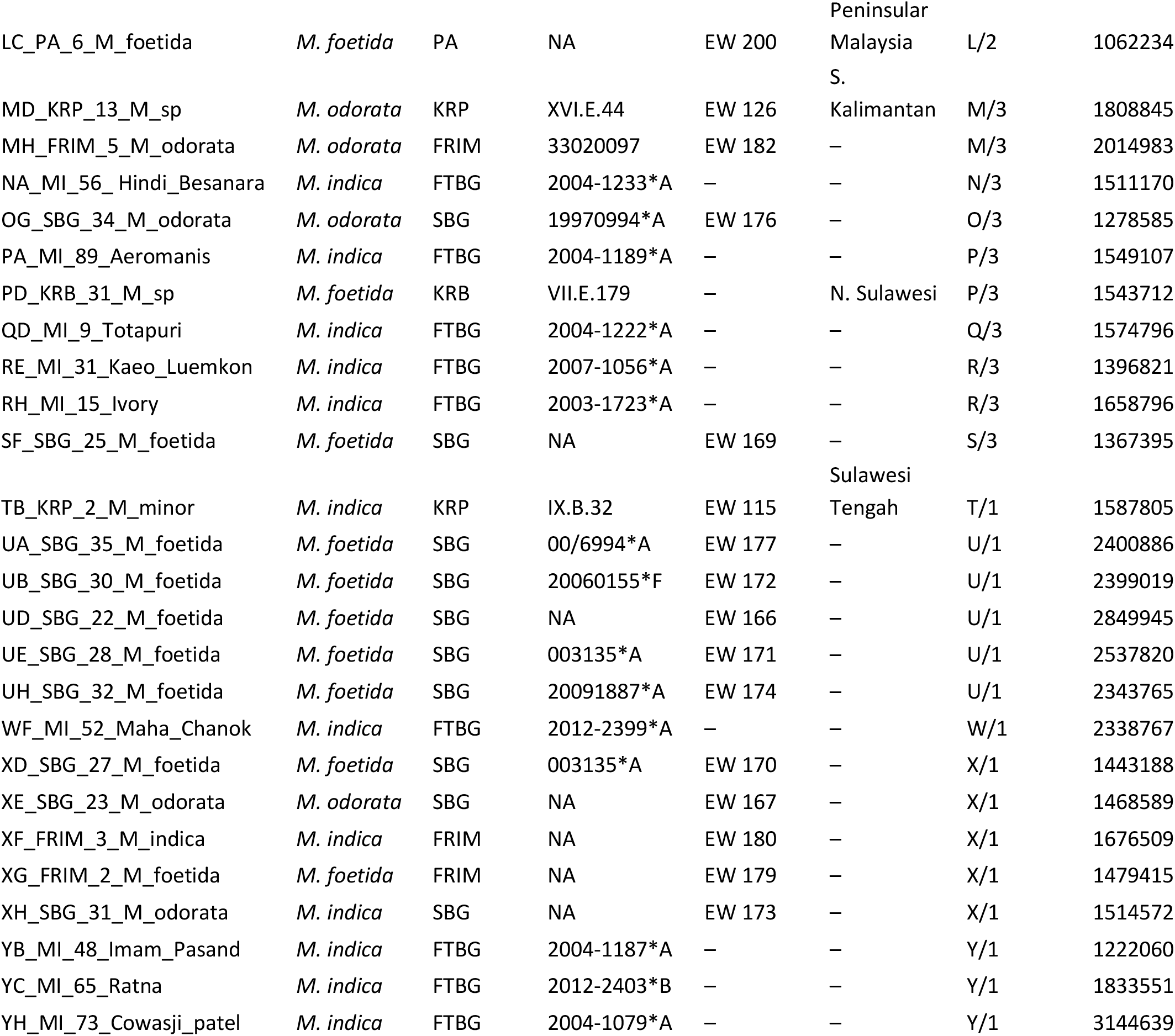
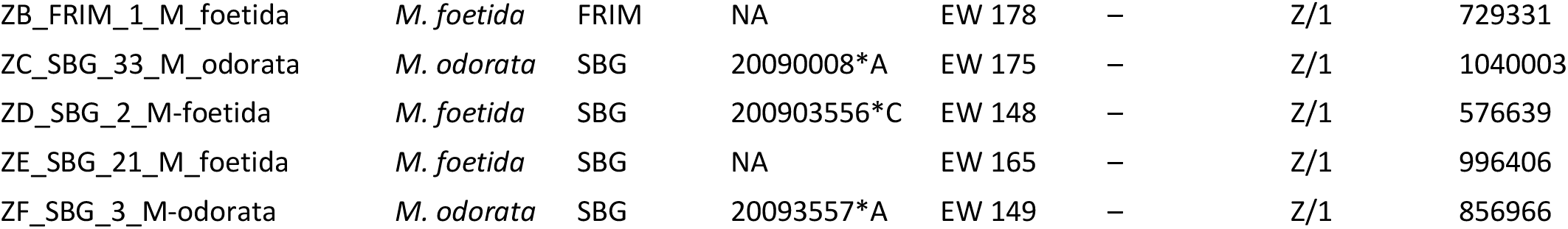
Samples in the M. odorata dataset. Sample ID consists of the individual ddRAD Sample ID, the study collection number and putative identification (cultivar or species name). The collection location (FTBG = Fairchild Tropical Botanic Garden, FTG = Fairchild Tropical Garden Herbarium, SBG = Singapore Botanic Garden, KRP = Purwodadi Botanic Garden, KRB = Bogor Botanic Garden, USDA = USDA Subtropical Horticulture Research Station at Chapman Field, FSP = Miami Dade Fruit and Spice Park, GBTB = Gardens by the Bay, FRIM = Forestry Research Institute of Malaysia, PA = Pasoh Forest Arboretum, PF = Pasoh Forest Research Station) and accession number within the respective collection are provided. The ddRAD library, sublibrary, and individual sample ID are given, as well as the number of raw reads for each individual.

**Table 2.**
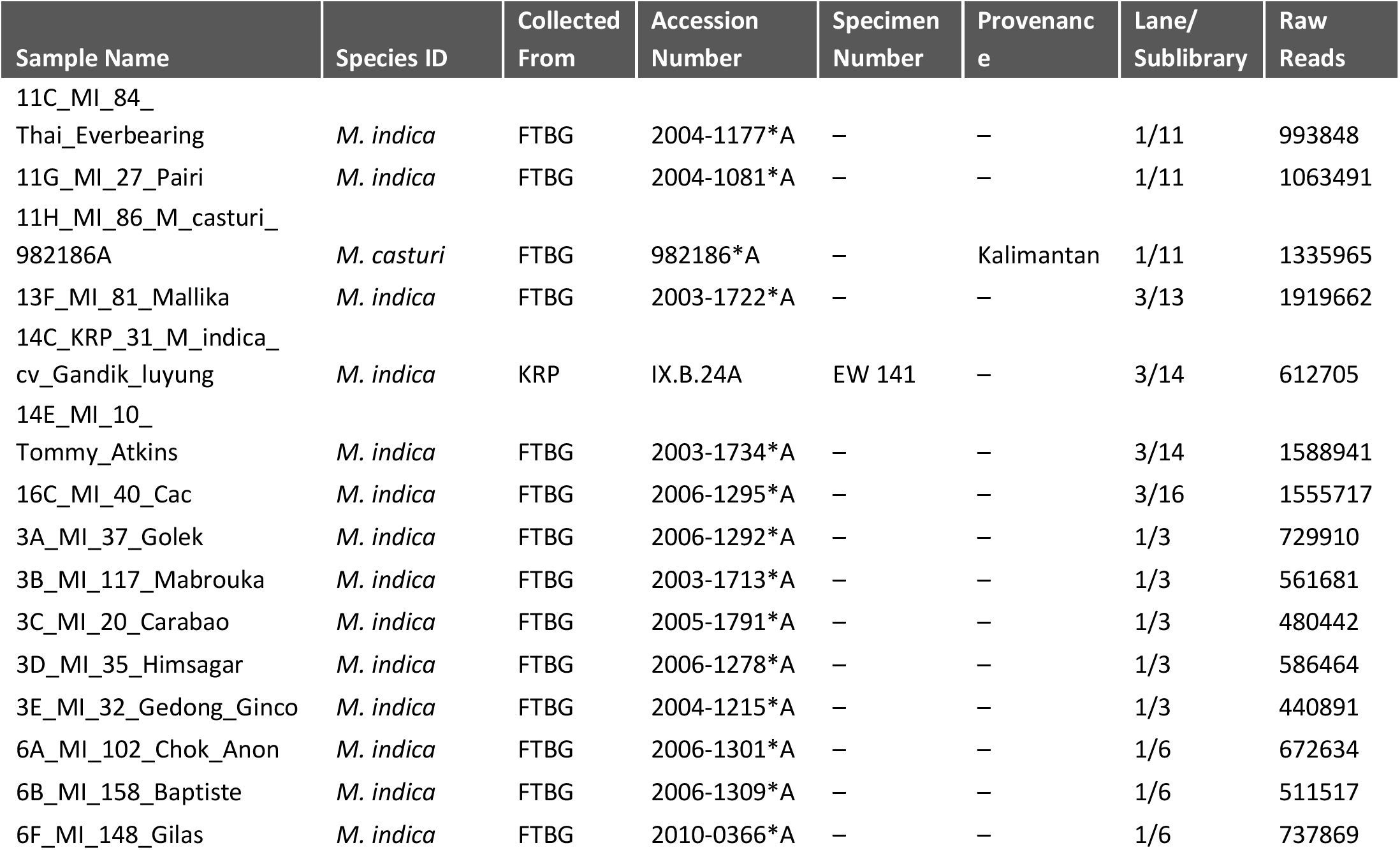
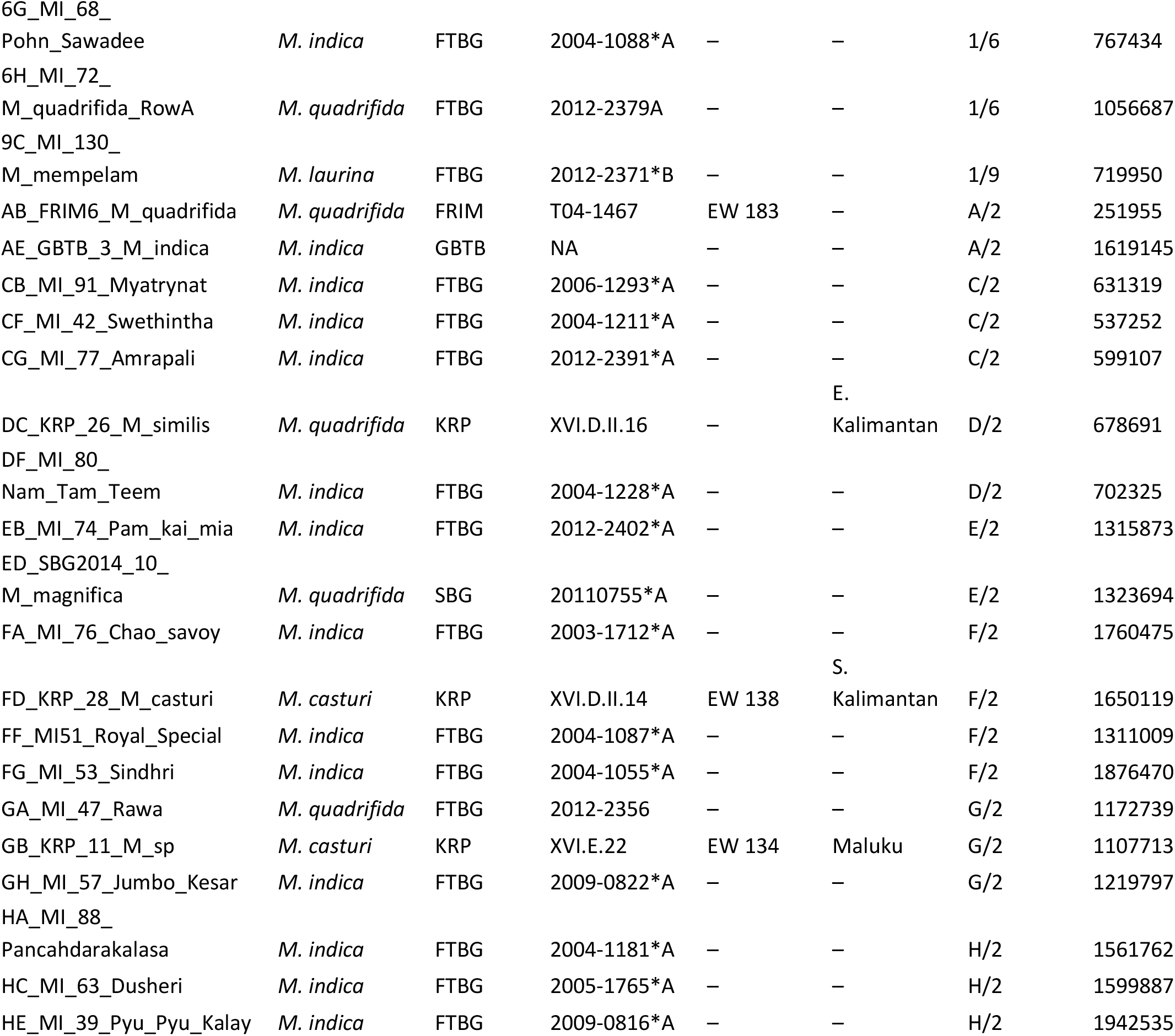
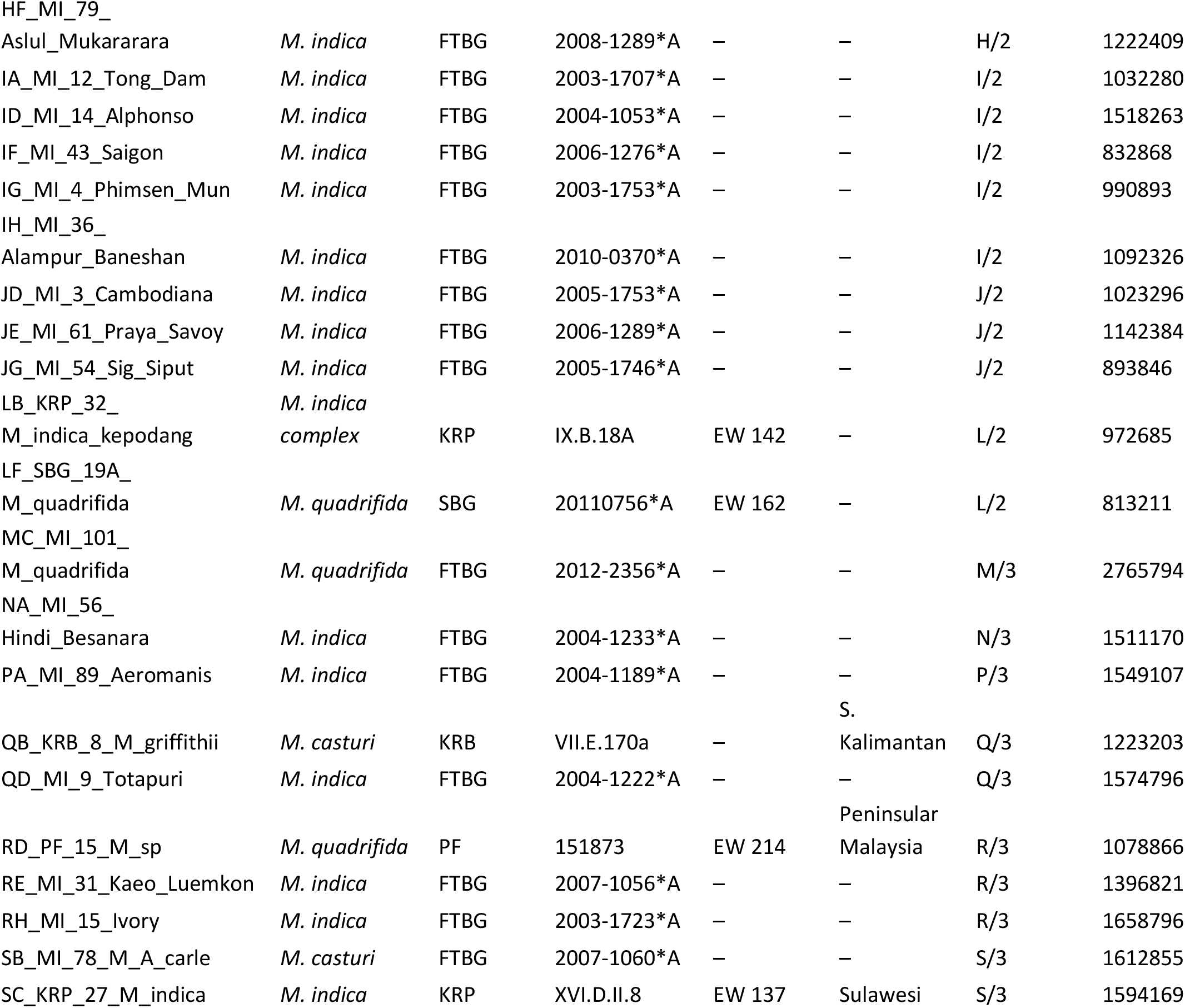
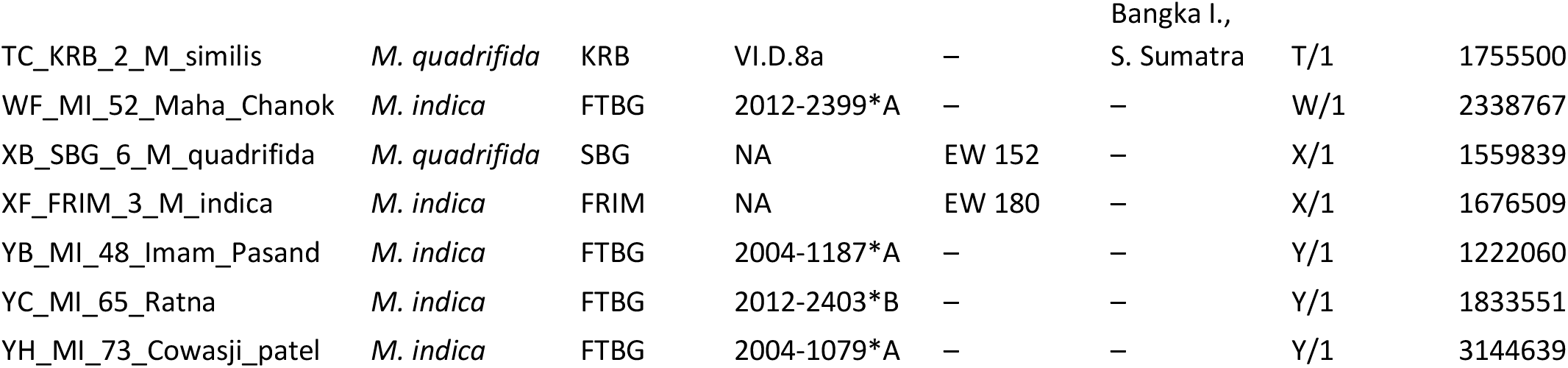
Samples in the M. casturi dataset. Sample ID consists of the individual ddRAD Sample ID, the study collection number and putative identification (cultivar or species name). The collection location (FTBG = Fairchild Tropical Botanic Garden, FTG = Fairchild Tropical Garden Herbarium, SBG = Singapore Botanic Garden, KRP = Purwodadi Botanic Garden, KRB = Bogor Botanic Garden, USDA = USDA Subtropical Horticulture Research Station at Chapman Field, FSP = Miami Dade Fruit and Spice Park, GBTB = Gardens by the Bay, FRIM = Forestry Research Institute of Malaysia, PA = Pasoh Forest Arboretum, PF = Pasoh Forest Research Station) and accession number within the respective collection are provided. The ddRAD library, sublibrary, and individual sample ID are given, as well as the number of raw reads for each individual.

### Library preparation

Double-digest restriction site-associated DNA sequencing (ddRADseq) libraries were prepared according to the protocol of Peterson et al. (2012). Briefly, 300-1000 ng genomic DNA was digested with NlaIII and MluCI before ligating on custom-designed adapters that contained one of eight unique barcode sequences. Groups of eight individuals with different barcode sequences were pooled together to form a sublibrary prior to size selection using a Pippin Prep (Sage Science) that targeted 350 bp inserts (tight size selection at 425 bp using an external marker). To amplify sublibraries and add a unique PCR index, short cycle PCR was performed, using six separate reactions to avoid PCR bias. The quality of amplified sublibraries was checked on an Agilent Bioanalyzer (DNA high sensitivity chip). For any sublibraries where overamplification occurred, size selection on Pippin Prep was performed again to remove non-target DNA. Following size selection, 12 sublibraries consisting of 96 individuals total were pooled in equimolar amounts. Each of the three libraries was sequenced at the University of Southern California’s Genome and Cytometry Core in a single lane of rapid run on an Illumina HiSeq 2500.

### Bioinformatics

Raw fastq files were quality checked using the software FASTQC v.0.11.4 (Andrews, 2010). The ipyRAD bioinformatic pipeline (Eaton, 2014) was used to analyze raw reads under default parameters except for the following: a single mismatched base was allowed within the barcode sequence, adapter filtering was set to option 2, maximum read depth for loci was set to 1000. The 280 individuals that were sequenced were subset into the *M. odorata* and *M. casturi* datasets (90 and 65 individuals, respectively) in the ipyRAD workflow. Clustering within and between individuals was performed at 85% identity. For this study, the minimum number of individuals required to have data at a given locus was set to filter out loci with >10% missing data for the *M. odorata* dataset and >50% missing data for the *M. casturi* dataset.

### Population structure and admixture

Population structure and admixture of the *M. odorata* and *M. casturi* datasets were analyzed independently using the Bayesian K-means program STRUCTURE v. 2.3.4 (Pritchard et al., 2000; Falush et al., 2003; Hubisz et al., 2009). First, an average value of lambda for K=1 was estimated across 10 runs of 100,000 iterations with 10,000 steps of burn-in. Population structure was estimated for K of 1 to 10, using the estimated value of lambda. For each value of K, 10 runs of 100,000 iterations with 10,000 steps of burn-in were completed. The optimal value of K was determined according to the deltaK parameter of Evanno et al. (2005) in the program STRUCTURE HARVESTER v. 0.6.94 (Earl and vonHoldt, 2012). For each value of K, the program CLUMPP v. 1.1.2 (Jakobsson and Rosenberg, 2007) was used to summarize results across the 10 runs performed using the greedy option (M=2) for K=1-7 or the LargeKGreedy option (M=3) for K=8na -10 with G’ similarity and 1,000 random permutations. Summarized results were visualized using the program DISTRUCT v. 1.1 (Rosenberg, 2004).

Population structure was also assessed independently for the *M. odorata* and *M. casturi* datasets using principal component analysis (PCA) in the R package adegenet (Jombart, 2008; Jombart and Ahmed, 2011). For the *M. odorata* dataset, two individuals identified as outliers were removed from PCA analysis and downstream analyses, resulting in a dataset of 88 individuals. The R package INTROGRESS (Gompert and Buerkle, 2010) was used to calculate hybrid indices (Buerkle, 2005) and interspecific heterozygosity for the *M. odorata* and *M. casturi* datasets. Additionally, a neighbor network analysis was performed on the *M. odorata* and *M. casturi* datasets using the program SPLITSTREE (Huson and Bryant, 2006) under default parameters.

### Population Genetics

Multiple indices of genetic diversity were calculated for hybrid and parental species in the *M. odorata* and *M. casturi* datasets. Percent polymorphism, observed heterozygosity, and gene diversity were calculated using the R package adegenet (Jombart, 2008; Jombart and Ahmed, 2011). Allelic richness and the number of private alleles (SNPs) in each species were calculated in the R packages PopGenReport (Adamack and Gruber, 2014) and poppr (Kamvar et al., 2014), respectively. The number of private alleles was also calculated as a percent of the total number of SNP loci analyzed within each dataset. For *M. odorata* and *M. casturi*, the index *rbarD* (Agapow and Burt, 2001), a standardized form the index of association (Brown et al., 1980) that assesses multilocus linkage disequilibrium, was calculated in poppr using 999 permutations to test for significance.

## Results

### Sequencing Results

For the entire 280 individuals sequenced across three lanes, we obtained 454,840,461 raw reads, with 116,110,642 for the 90 individuals included in the *M. odorata* dataset (average: 1,290.118, SD: 558,342) and 80,460,503 for the 65 individuals in the *M. casturi* dataset (average: 1,237,854, SD: 550,028). FastQC results indicated reads were of high quality across the entire 150 bp length. After filtering for missing data, we recovered 481 unlinked SNPs from the *M. odorata* dataset and 3,609 unlinked SNPs from the *M. casturi* dataset.

### Mangifera odorata

For the *M. odorata* dataset, analysis of population structure in the Bayesian software STRUCTURE found *K* = 2 to be the optimal number of populations according to the Δ*K* method of Evanno (Fig. 1, Fig. S1). All samples of the parental species *M. foetida* were assigned to a single population, with high levels of identity (99.1-100.0%). The Indian and Southeast Asian populations of *M. indica* were assigned to a second population, with individual identity ranging from 99.9-100%. No differentiation was observed between Indian and Southeast Asian populations for *K =* 2. All 17 individuals of *M. odorata* showed high levels of admixture, with 58.73-74.01% of ancestry assigned to the *M. foetida* population and 25.99-41.27% of ancestry assigned to the *M. indica* population. Because the Δ*K* method of Evanno is known to identify only the highest existing level of population structure, larger values of *K* were examined, but were not found to show any additional meaningful structure.

**Figure 1.**
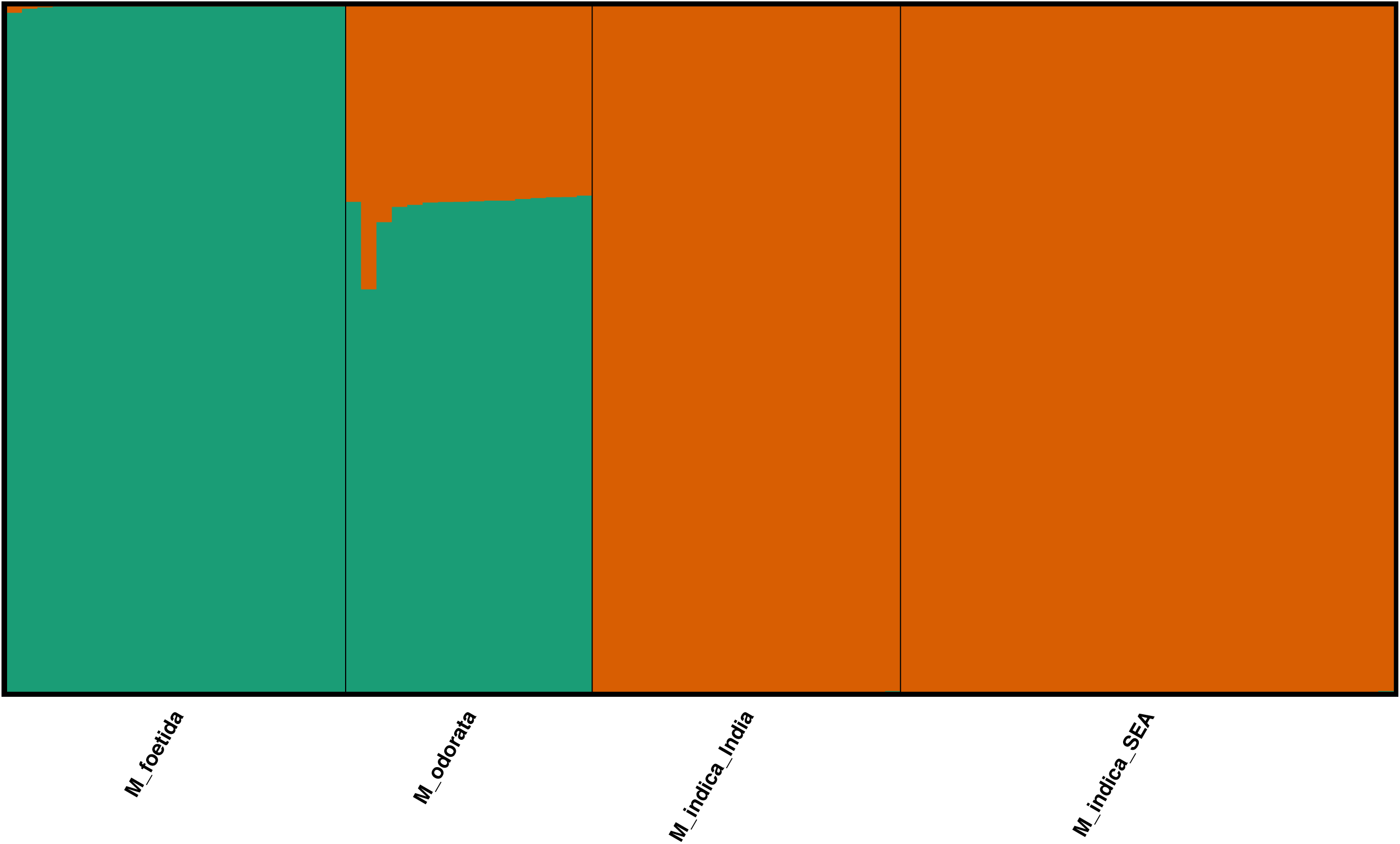
Visualization of inferred population structure for the *M. odorata* dataset produced using the software distruct with *K =* 2 populations. Individuals within the dataset are represented by a vertical bar and colors represent assignment to each population. Individuals are labeled by species name *(M. foetida, M. odorata, M. indica)* and, for *M. indica*, region of origin (India, Southeast Asia [SEA]).

Analysis of the *M. odorata* dataset using principal components shows three clusters, which correspond to *M. indica, M. odorata*, and *M. foetida* (Fig. 2). The first principal component (PC1) accounts for 37.68% of the variation observed in the data and separates the three species, with *M. odorata* recovered as intermediate to *M. foetida* and *M. indica*. The second principal component (PC2) accounts for 6.32% of the variation present in the dataset and further distinguishes *M. odorata* from *M. foetida* while also stratifying Southeast Asian and Indian populations of *M. indica*.

**Figure 2.**
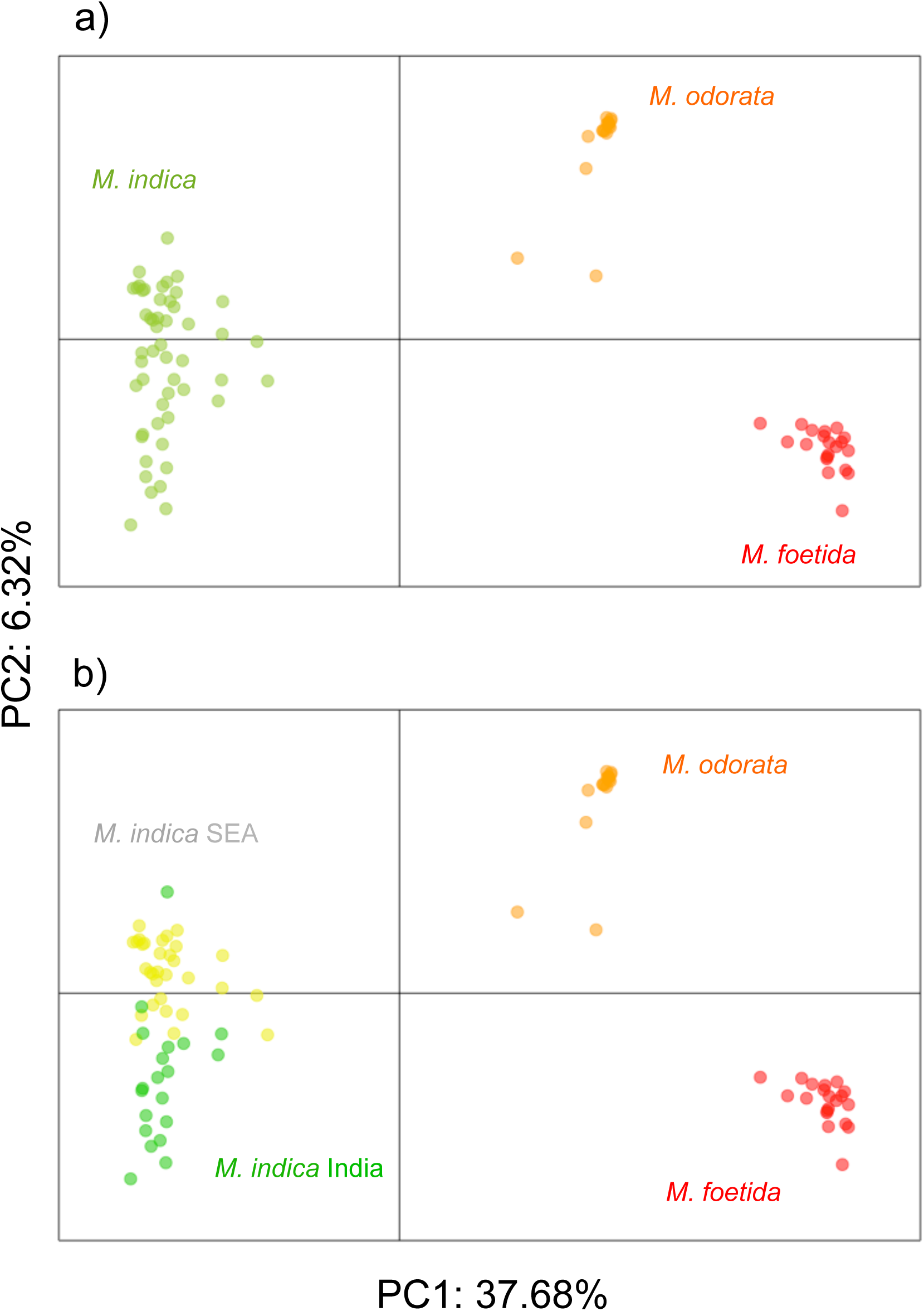
Principal component analysis of 88 samples of *M. foetida, M. odorata*, and *M. indica* from the *M. odorata* dataset, with individual samples colored according to a) species only (red = *M. foetida*, orange = *M. odorata*, green = *M. indica*) or b) species and region (red = *M. foetida*, orange = *M. odorata*, green = Indian *M. indica*, yellow = Southeast Asian *M. indica*).

Of the 17 *M. odorata* individuals examined, 16 have a very similar hybrid index (0.32- 0.36, where 0 is *M. foetida* and 1 is *M. indica*) and similar values of interspecific heterozygosity (23.29-25.51%) (Fig. 3). The remaining individual is genetically distinct, with an estimated hybrid index of 0.44 and interspecific heterozygosity of 17.86%. Network analysis of the *M. odorata* dataset places all samples of *M. odorata* directly on the branch separating *M. indica* and *M. foetida*, though the branch length between *M. odorata* and *M. foetida* is shorter than that between *M. odorata* and *M. indica* (Fig. 4).

**Figure 3.**
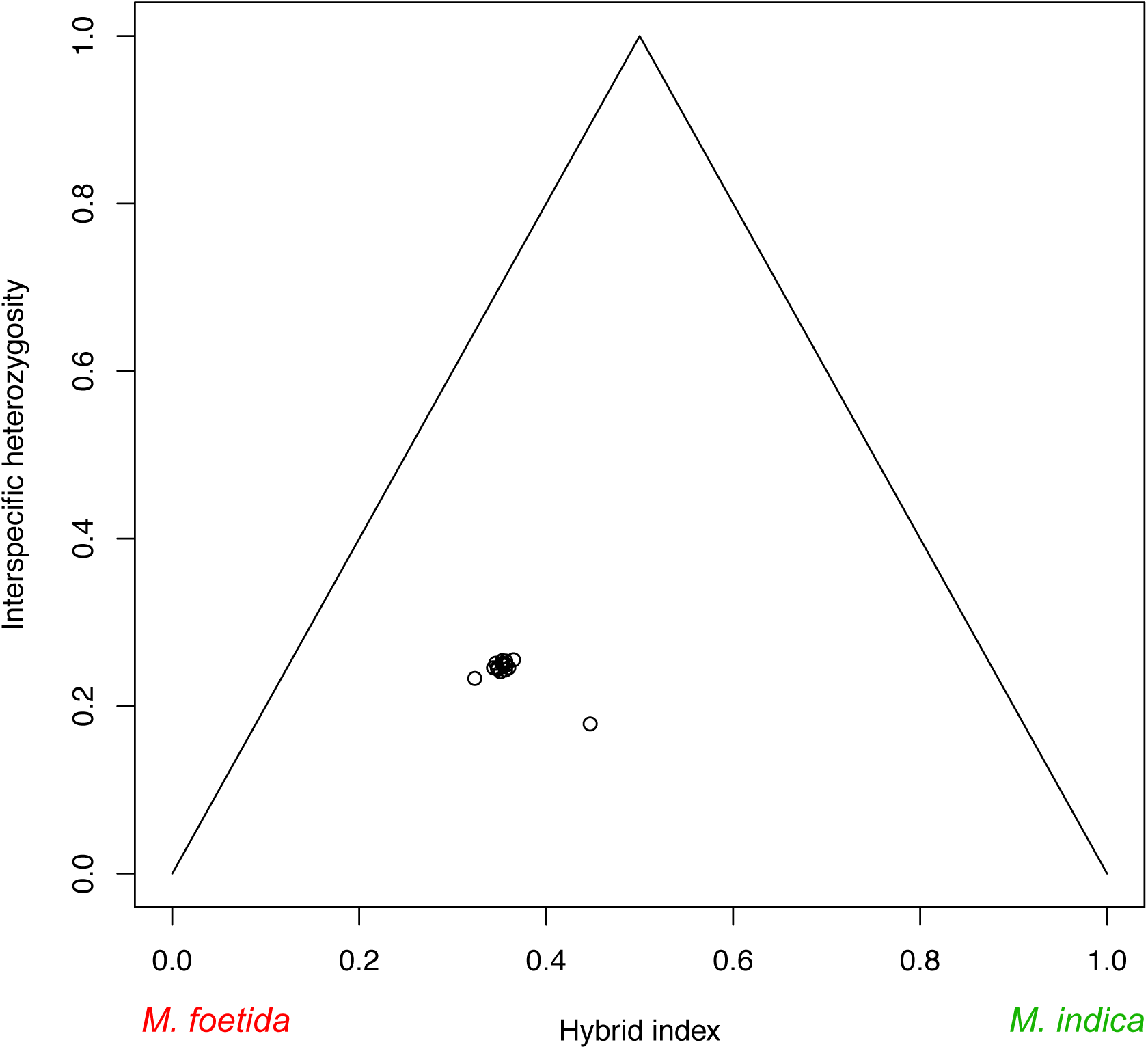
Plot of hybrid index (0 = *M. foetida*, 1 = *M. indica*) and interspecific heterozygosity for 17 samples of *M. odorata* calculated in the program INTROGRESS.

**Figure 4.**
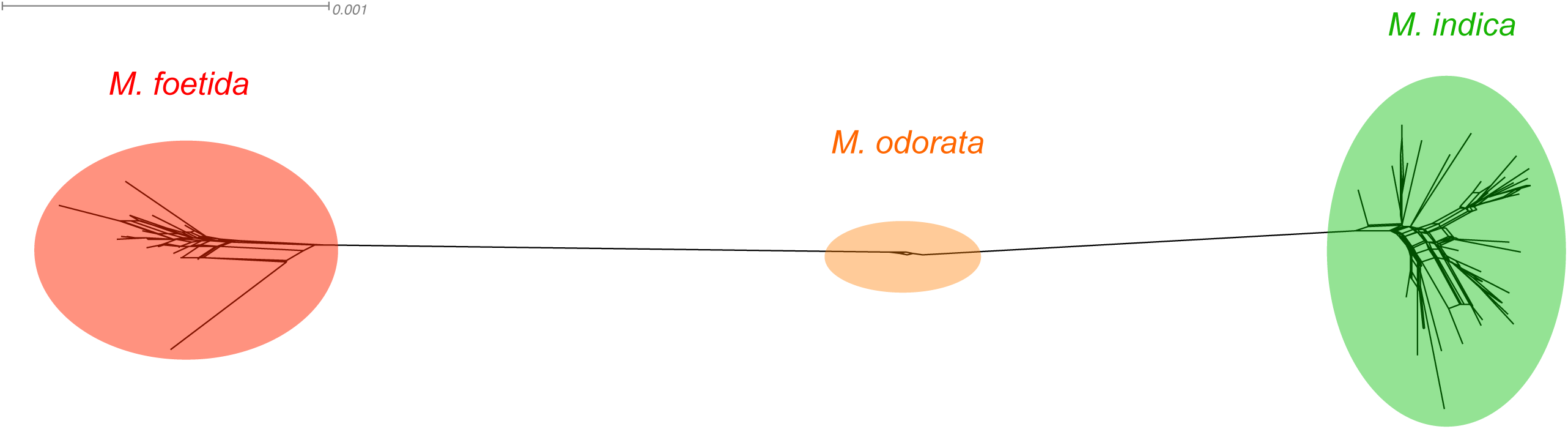
Neighbor network tree for *M. odorata* (orange)*, M. foetida* (red), and *M. indica* (green) inferred using SPLITSTREE.

Indices of genetic diversity for the *M. odorata* dataset (Table 3) show that the hybrid individuals have higher levels of observed heterozygosity (*H_O_* = 0.3005) than *M. indica* (*H_O_* = 0.1072) and *M. foetida* (*H_O_* = 0. 0487). Gene diversity is highest in *M. odorata (H_S_* = 0.1638), followed by *M. indica* (*H_S_* = 0.1186) and *M. foetida (H_S_* = 0.0644). A negative inbreeding coefficient for *M. odorata (F_IS_* = -0.6230) indicates an excess of outcrossing, while *M. indica* and *M. foetida* have values indicating slight inbreeding *(F_IS_* = 0.0840 and 0.3115, respectively). *Mangifera indica* has the lowest percent polymorphism (25.32%), while *M. foetida* has the highest (57.29%) and intermediate levels are seen in *M. odorata* (43.22%). Similar values of nucleotide diversity are observed in *M. indica* and *M. foetida* (0.0309 and 0.0246, respectively) while *M. odorata* has very low nucleotide diversity (0.0016). For *M. odorata*, values of *rbarD*, a measure of multilocus linkage disequilibrium, are significantly different from zero (*rbarD* = 0.3984, *p =* 0.001, Fig. S2). The number and percent of private alleles across all loci is highest for *M. indica* (173, 44.25%), lower in *M. foetida* (46, 11.76%), and very low in *M. odorata* (1, 0.26%).

**Table 3.** . Population genetic analysis of *M. odorata* and its parental taxa, *M. indica* and *M. foetida. Ho*: observed heterozygosity; *Hs*: heterozygosity within populations, aka ‘gene diversity’; *Fis*: inbreeding coefficient; *π:* nucleotide diversity; rbarD: multilocus linkage disequilibrium; Ap%: percent private alleles.

| Species | Ho | Hs | Fis | %poly | $\pi$ | rbarD | Ap% |
| --- | --- | --- | --- | --- | --- | --- | --- |
| *M. odorata* | 0.3005 | 0.1638 | -0.6230 | 43.22% | 0.0016 | 0.3984 | 0.26% |
| *M. indica* | 0.1072 | 0.1186 | 0.0840 | 25.32% | 0.0309 | 0.0270* | 44.25% |
| *M. foetida* | 0.0487 | 0.0644 | 0.3115 | 57.29% | 0.0246 | 0.0407 | 11.76% |

### Mangifera casturi

Analysis of the *M. casturi* dataset in the program STRUCTURE found *K* = 3, 5, and 8 to be equally optimal according to the Δ*K* method of Evanno (Fig. 5, Fig. S3). Patterns of genetic structure are generally similar across the three values of *K.* For *K* = 3, *M. indica* samples are assigned high identity to a single population (78.38-84.63%) with moderate identity from a secondary population (13.43-16.92%) and low levels of identity from a third population (3.06- 5.47%). Samples from *M. quadrifida* show a distinct pattern, with highest identity coming from the third population (50.82-60.44%), and moderate levels from both the second (18.00-21.50%) and first (18.05-30.42%) populations. The samples of *M. casturi* are intermediate to the putative parental populations: assignment to the first population ranges from 58.02-59.77%, with assignment to the second population at 18.57-19.83% and assignment to the third population at 21.63-22.39%. As the number of populations increases from three to five to eight, the population assignments for *M. indica* and *M. quadrifida* become more distinct, while *M. casturi* continues to have assignments that are intermediate to the putative parental species and shows no unique ancestry.

**Figure 5.**
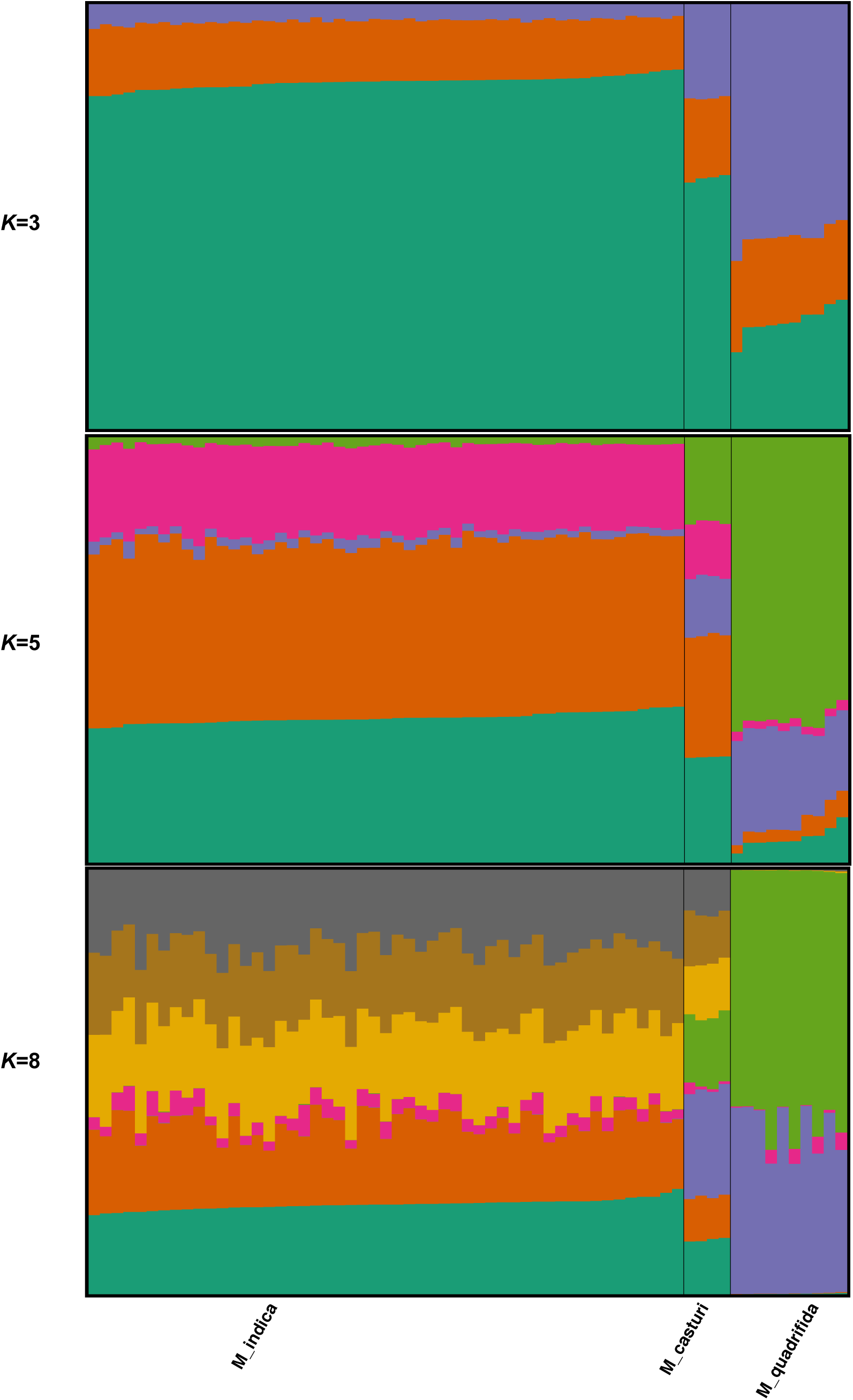
Visualization of inferred population structure for the *M. casturi* dataset produced using the software distruct with *K =* 2 populations. Individuals within the dataset are represented by a vertical bar and colors represent assignment to each population. Individuals are labeled by species name *(M. quadrifida, M. casturi, M. indica)*.

For the *M. casturi* dataset, principal component analysis recovers four clusters of individuals, with one cluster corresponding to *M. indica*, a second corresponding to *M. casturi*, and two clusters representing *M. quadrifida* (Fig. 6). The first principal component accounts for 27.20% of the variation in the dataset and separates *M. casturi* and *M. quadrifida* from *M. indica*. The second principal component accounts for 9.73% of the variation in the dataset and separates *M. casturi* and the two populations of *M. quadrifida*.

**Figure 6.**
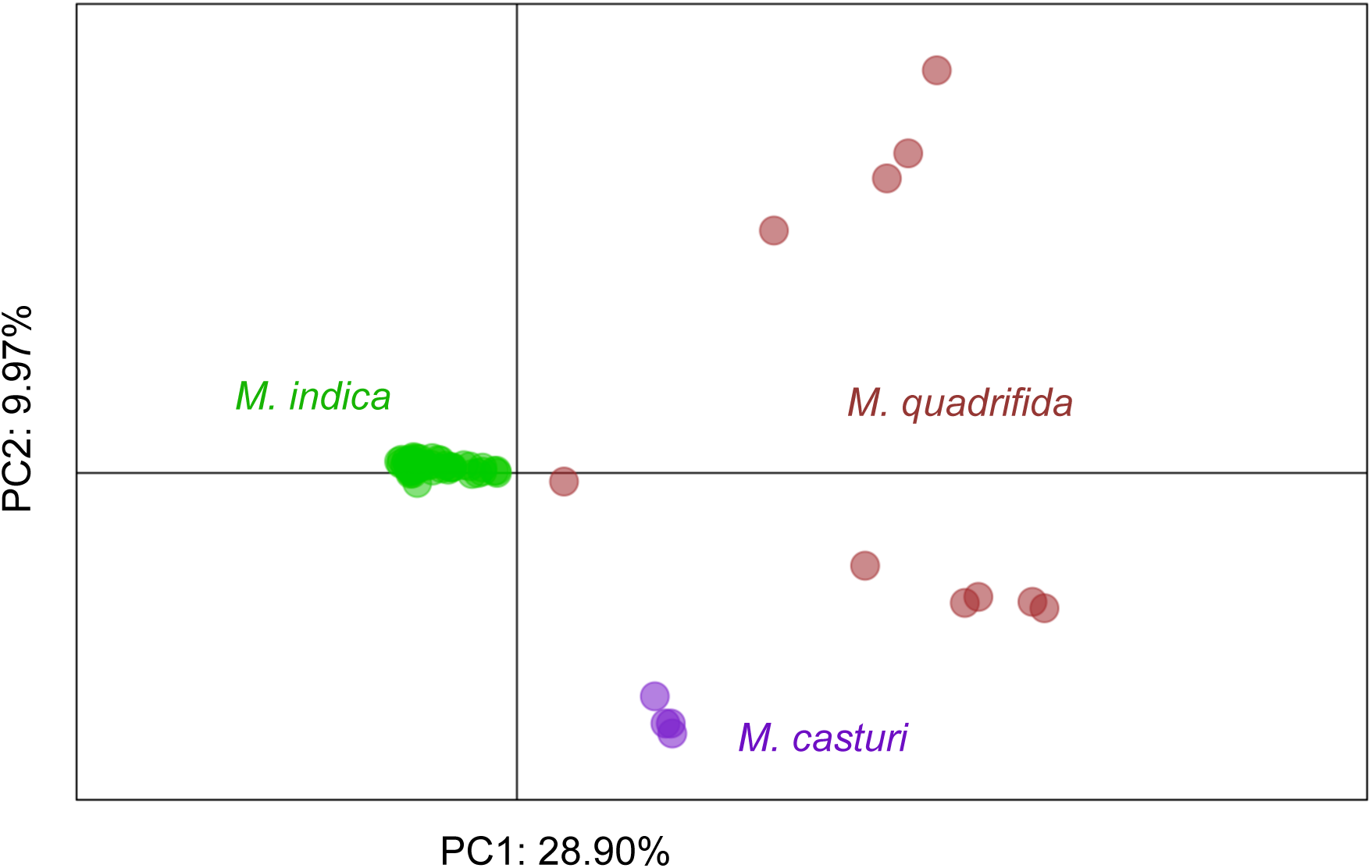
Principal component analysis of 65 samples of *M. quadrifida, M. casturi*, and *M. indica* from the *M. casturi* dataset, with individual samples colored according to a) species only (brown = *M. quadrifida*, purple = *M. casturi*, green = *M. indica*).

Analysis of the four samples of *M. casturi* using the INTROGRESS software indicates that the samples are very similar, with hybrid indices (where 0 is *M. quadrifida* and 1 is *M. indica*) between 0.60-0.62 and interspecific heterozygosity of 25.3-26.3% (Fig. 7). Network analysis of the *M. casturi* dataset in the program SPLITSTREE places *M. casturi* directly intermediate to *M. indica* and *M. quadrifida*. However, the neighbor network identifies two populations of *M. quadrifida*, one of which is also found to be an intermediate (Fig. 8). As a result of the population differentiation of *M. quadrifida* observed in the PCA and SPLITSTREE analyses, we attempted to analyze the *M. casturi* dataset independently for each of the *M. quadrifida* populations, but the small sample sizes of the individual *M. quadrifida* populations prohibited robust analysis.

**Figure 7.**
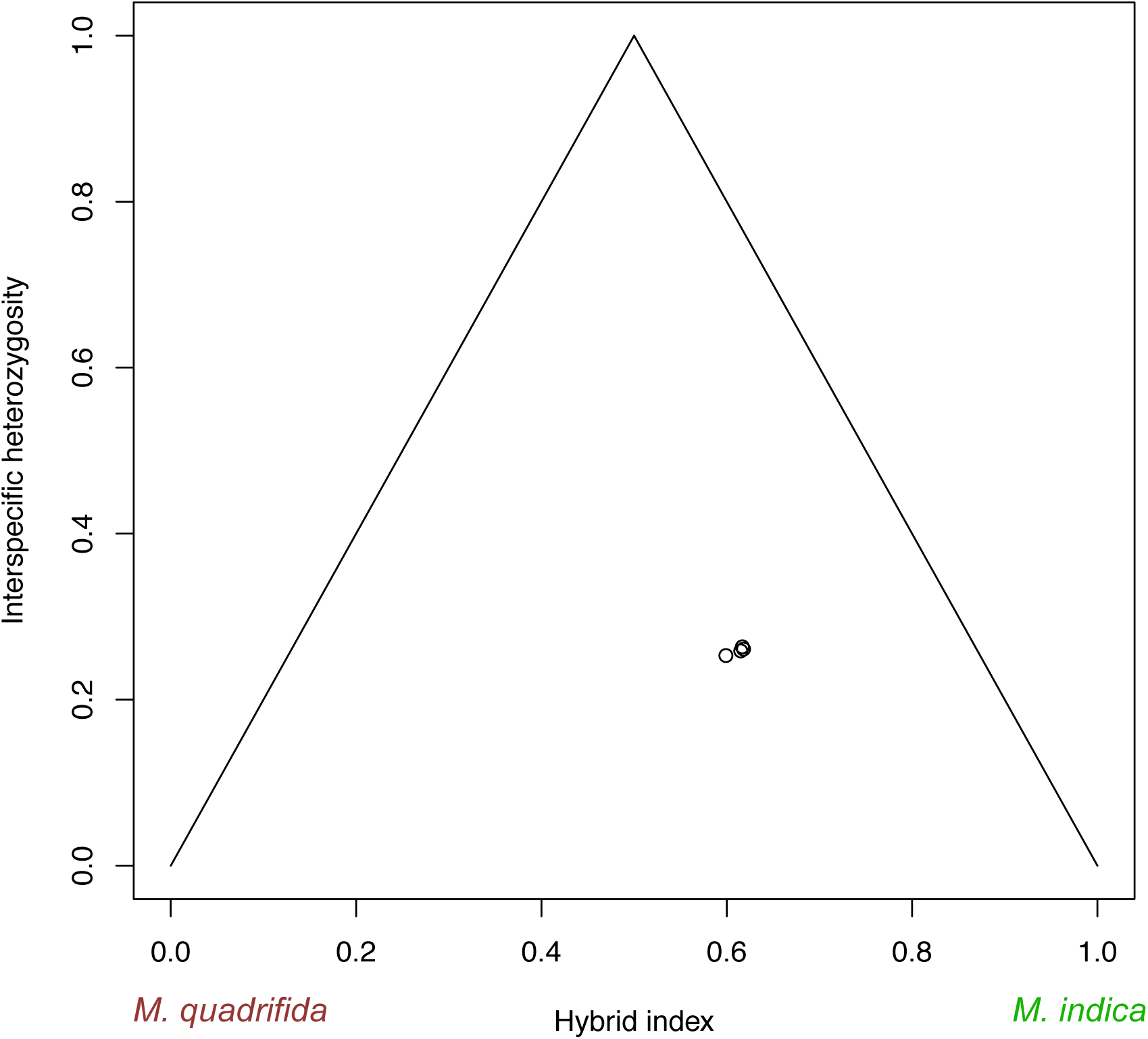
Plot of hybrid index (0 = *M. quadrifida*, 1 = *M. indica*) and interspecific heterozygosity for 4 samples of *M. casturi* calculated in the program INTROGRESS.

**Figure 8.**
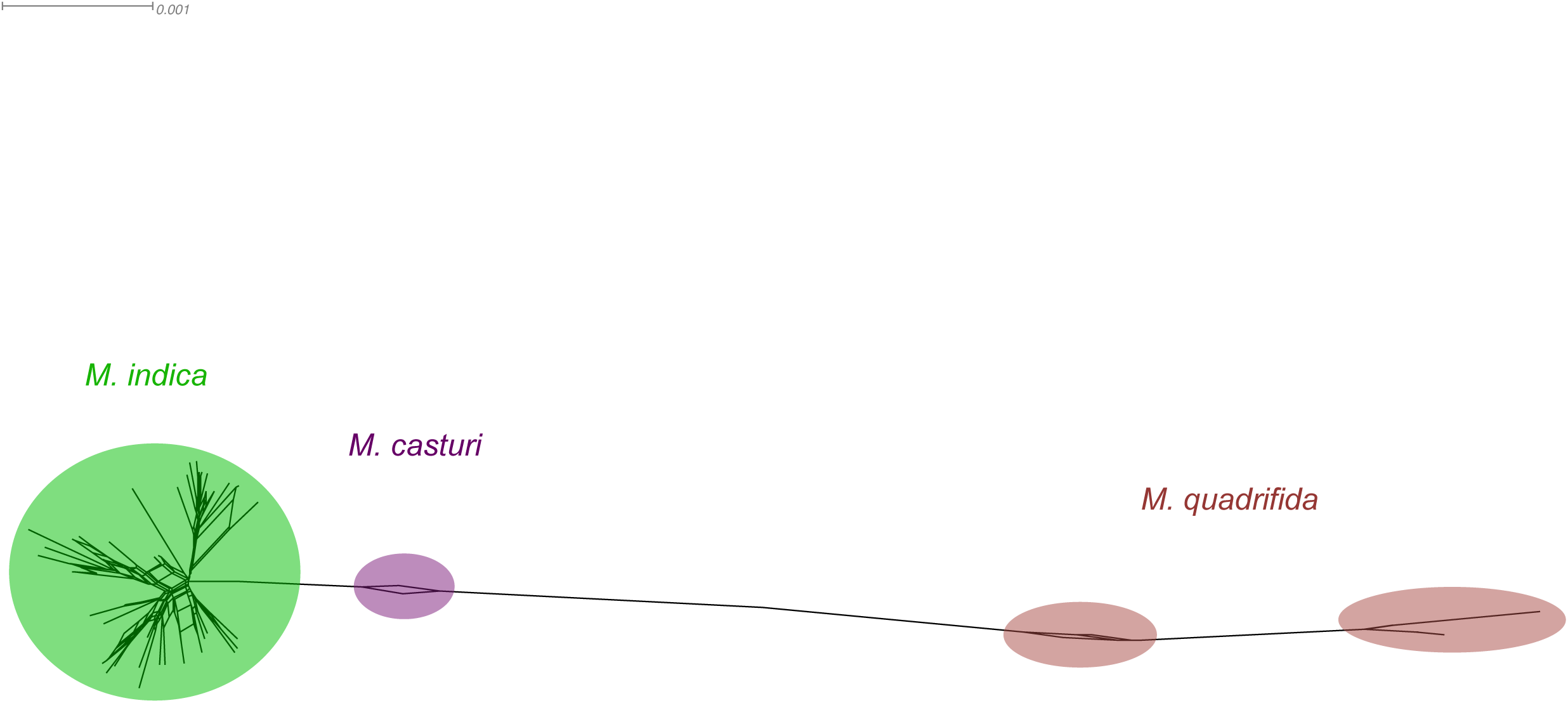
Neighbor network tree for *M. casturi* (purple)*, M. quadrifida* (brown), and *M. indica* (green) inferred using SPLITSTREE.

Genetic diversity indices of the *M. casturi* dataset (Table 4) show similar patterns to those observed in the *M. odorata* dataset. Individuals of *M. casturi* have higher observed heterozygosity (*H_O_* = 0. 2869) than the putative parental taxa (*M. indica H_O_* = 0.0939, *M. foetida H_O_* = 0.1473). Gene diversity was lowest in *M. indica (H_S_* = 0.1112), and similar in *M. casturi* and *M. quadrifida (H_S_* = 0.1580 and 0.1377, respectively). A strongly negative inbreeding coefficient for *M. casturi (F_IS_* = -0.7801) indicates an excess of outcrossing in this species, while values closer to zero were calculated for *M. indica* and *M. quadrifida (F_IS_* = 0.1213 and -0.0394, respectively). The highest percent polymorphism is observed in *M. indica* (53.75%), while *M. quadrifida* and *M. casturi* have similar values (37.60% and 32.70%, respectively). Similar values of nucleotide diversity are observed in *M. indica* and *M. quadrifida* (0.0321 and 0.0331, respectively) while the value for *M. casturi* is much lower (0.0045). For both *M. odorata* and *M. quadrifida*, values of *rbarD*, a measure of multilocus linkage disequilibrium, are significantly different from zero (*rbarD* = 0.2522 and 0.3011, *p =* 0.001, Fig. S4). The number and percent of private alleles across all loci is highest for *M. indica* (1453, 40.26%), lower in *M. quadrifida* (937, 25.96%), and lowest in *M. casturi* (62, 1.72%).

**Table 4.** Population genetic analysis of M. casturi and its putative parental taxa, M. indica and M. quadrifida. Ho: observed heterozygosity; Hs: heterozygosity within populations, aka ‘gene diversity’; Fis: inbreeding coefficient; π: nucleotide diversity; rbarD: multilocus linkage disequilibrium; Ap%: percent private alleles.

| Species | Ho | Hs | Fis | %poly | $\pi$ | rbarD | Ap% |
| --- | --- | --- | --- | --- | --- | --- | --- |
| *M. casturi* | 0.2869 | 0.1580 | -0.7801 | 32.70% | 0.0045 | 0.2522 | 1.72% |
| *M. indica* | 0.0939 | 0.1112 | 0.1213 | 53.75% | 0.0321 | 0.0455 | 40.26% |
| *M. quadrifida* | 0.1473 | 0.1377 | -0.0394 | 37.60% | 0.0331 | 0.3011 | 25.96% |

## Discussion

In this study, we use RADseq to explore the consequences of hybridization between a widely cultivated fruit tree, *M. indica*, and two different congeners in Southeast Asia. We find support for the previously demonstrated hybrid origin of *M. odorata* and, for the first time, evidence that *M. casturi* is also of hybrid origin. Both *M. odorata* and *M. casturi* lack unique genetic diversity compared to their parental taxa, as indicated by the low number of private alleles present in hybrid populations and by population structure and network analyses. However, neither *M. odorata* nor *M. casturi* is directly intermediate to its respective parental taxa, as would be expected for first generation (F1) hybrids. Instead, both hybrids have levels of interspecific heterozygosity near 25%, indicating that they are likely the result of a backcross with one parental species. In the case of *M. odorata*, our data support the results of Teo et al. (2002), who found it to be more closely related to *M. foetida*. Therefore, *M. odorata* is probably the product of a backcross between an F1 hybrid and *M. foetida.* On the other hand, *M. casturi* has greater genetic affinity to *M. indica* than to *M. quadrifida*, and is most likely the result of an F1 hybrid backcross with *M. indica*.

Although *M. odorata* is of hybrid origin, we find no evidence of introgression between *M. indica* and *M. foetida* via the hybrid intermediate *M. odorata.* On the contrary, the genetic similarity of 16 of the 17 individuals of *M. odorata*, as evidenced by significant multilocus linkage disequilibrium and similar measures of admixture (interspecific heterozygosity, hybrid index, population structure assignment) combined with strongly negative inbreeding coefficients suggest that *M. odorata* is the result of a limited number of hybridization events. A previous study of diversity in 11 landraces of *M. odorata* using microsatellite markers also found it to have a very narrow genetic base (Yamanaka et al., 2006). We find patterns of diversity in *M. casturi* to be similar to *M. odorata*, with significant multilocus linkage disequilibrium, and individuals having very similar measures of admixture. Notably, in the case of *M. casturi* we find some evidence indicating population differentiation within *M. quadrifida* that may be the result of alternate backcrossing events (e.g., an F1 hybrid backcross with *M. quadrifida)*.

The remarkably similar genetic identity of individuals within *M. odorata* and *M. casturi* is consistent with hybrids that have formed only a few times and have thereafter been maintained clonally. In most cases of fruit trees, grafting is the primary means of clonal propagation of tree crops, allowing unique genetic individuals to be maintained in perpetuity (e.g., Warschefsky et al., 2016). Considering the small number of *M. casturi* individuals analyzed, it is possible that the four individuals we included were grafted clones. However, in the case of *M. odorata*, our 17 samples originated from different sources and the majority of individuals were not apparently grafted. So, if not by grafting, how could hybrid *M. odorata* be clonally maintained?

Nucellar polyembryony, a type of asexual reproduction or apomixis, produces multiple embryos within a single seed. One of the embryos in a polyembryonic seed is sexually derived, while the others develop from the maternal nucellar tissue (Aleza et al., 2010). While rare in angiosperms, nucellar polyembryony is well documented in many *Citrus* species (Wang et al., 2017) along with Southeast Asian cultivars of *M. indica* (Mukherjee and Litz, 2009), and has been reported at least three other species of *Mangifera*, including *M. odorata* and *M. casturi* (Kostermans and Bompard, 1993; Mukherjee and Litz, 2009; Lim, 2012a; b). Therefore, we propose that *M. odorata* may represent a cultivated hybrid lineage maintained by clonal reproduction via nucellar polyembryony. We can also speculate that *M. casturi* represents a similar cultivated, polyembryonic hybrid lineage, though additional individuals should be analyzed to confirm the genetic uniformity of the species.

Polyembryony is an important agricultural characteristic for tree crops like mango and citrus, allowing for the propagation of otherwise unattainable clonal rootstock material. Recent genome sequencing of multiple *Citrus* species has helped to shed light on the evolution of polyembryony in the group (Wang et al., 2017). In the case of citrus, the majority of cultivated species are of interspecific hybrid origin, (e.g., sweet orange, grapefruit), and it has been suggested that these hybrid lineages have been maintained by polyembryony, with different citrus cultivars originating through somatic mutations in clonal lineages rather than through sexual reproduction (Wu et al., 2018). Evidence from whole genome sequencing of multiple *Citrus* species indicates that nucellar polyembryony is controlled by a single dominant allele that first arose in the mandarin, *C. reticulata* (Wang et al., 2017).

In mango, traditional analysis of phenotypic segregation from crosses between polyembryonic and monoembryonic cultivars also ascertained that a single dominant gene controls the trait (Aron et al., 1998; Kuhn et al., 2017). However, at this point many questions about the process of nucellar polyembryony in *Mangifera* remain unanswered. One issue central to the aims of our study is the fate of the sexually reproduced embryos, both in the case of original F1 hybrids and of further backcrosses. Since we found no evidence of recurrent backcrossing or introgression, it is possible that these individuals do not commonly survive because of some genetic incompatibility. However, the lack of F1 individuals could be explained by nomenclature, as it is possible that *M. odorata* and *M. casturi* are names applied to only very specific hybrid lineages, much like the use of cultivar names, while F1 hybrids may be given different common and/or scientific names.

### Future Research

Here, we have provided preliminary evidence that indicates *M. casturi*, a species only known from cultivation and classified as extinct in the wild (IUCN, 2012), may be a hybrid of *M. indica* and *M. quadrifida.* More thorough sampling of *M. casturi* and *M. quadrifida* would provide greater insight into the origin of *M. casturi* and the population structure we observed within *M. quadrifida.* Although we analyzed more samples of *M. odorata* than *M. casturi*, the perplexing lack of diversity within *M. odorata* warrants additional sampling and investigation, particularly because the species was previously described as being polymorphic (Hou, 1978; Teo et al., 2002).

One important avenue for research is to determine whether *M. odorata* and *M. casturi* can be re-created by controlled crosses between their respective parental taxa. However, hand pollination of *Mangifera* species is said to be very difficult and is inefficient because a large proportion of fruits are aborted prematurely (Iyer and Schnell, 2009). In addition, given that *M. odorata* and *M. indica* appear to be backcrosses, replicating these individuals would require at least two generations, or 6-20 years (Iyer and Schnell, 2009).

Overall, our research provides important insights into the consequence of hybridization within *Mangifera*. Coupled with our knowledge of hybrid citrus species, our findings reveal a pattern of perennial crop cultivation: maintenance of favorable hybrid perennial crop lineages through apomixis.

## Notes

### Competing Interest Statement

The authors have declared no competing interest.

## Literature Cited

Abbott, R., D. Albach, S. Ansell, J.W. Arntzen, S.J.E. Baird, N. Bierne, J. Boughman, et al. 2013. Hybridization and speciation. Journal of Evolutionary Biology 26: 229–246.

Adamack, A.T., and B. Gruber. 2014. PopGenReport: Simplifying basic population genetic analyses in R. Methods in Ecology and Evolution 5: 384–387.

Agapow, P.M., and A. Burt. 2001. Indices of multilocus linkage disequilibrium. Molecular Ecology Notes 1: 101–102.

Aleza, P., J. Juárez, P. Ollitrault, and L. Navarro. 2010. Polyembryony in non-apomictic citrus genotypes. Annals of Botany 106: 533–545.

Anderson, E. 1949. Introgressive hybridization. John Wiley & Sons, Inc., New York, NY.

Anderson, E., and G.L. Stebbins. 1954. Hybridization as an evolutionary stimulus. Evolution 8: 378–388.

Andrews. S. 2010. FastQC: a quality control tool for high throughput sequence data. Available from: http://www.bioinformatics.babraham.ac.uk/projects/fastqc/.

Arnold, M. 1992. Natural hybridization as an evolutionary process. Annual Review of Ecology and Systematics 23: 237–261.

Arnold, M.L., E.S. Ballerini, and A.N. Brothers. 2012. Hybrid fitness, adaptation and evolutionary diversification: lessons learned from Louisiana Irises. Heredity 108: 159–66.

Aron, Y., H. Czosnek, and S. Gazit. 1998. Polyembryony in mango (*Mangifera indica* L.) is controlled by a single dominant gene. HortScience 33: 1241–1242.

Barton, N.H., and H. G.M. 1985. Analysis of hybrid zones. Annual Review of Ecology and Systematics 16: 113–148.

Bompard, J.M. 2009. Taxonomy and Systematics. In R. E. Litz [ed.], The mango: Botany, production and uses, CAB International, Cambridge, MA.

Brown, A.H., M.W. Feldman, and E. Nevo. 1980. Multilocus Structure of Natural Populations of *Hordeum spontaneum*. Genetics 96: 523–536.

Buerkle, C.A. 2005. Maximum-likelihood estimation of a hybrid index based on molecular markers. Molecular Ecology Notes 5: 684–687.

Coart, E., S. Van Glabeke, M. De Loose, A.S. Larsen, and I. Roldán-Ruiz. 2006. Chloroplast diversity in the genus *Malus*: new insights into the relationship between the European wild apple (*Malus sylvestris* (L.) Mill.) and the domesticated apple (*Malus domestica* Borkh.). Molecular ecology 15: 2171–82.

Cornille, A., P. Gladieux, M.J.M. Smulders, I. Roldán-Ruiz, F. Laurens, B. Le Cam, A. Nersesyan, et al. 2012. New insight into the history of domesticated apple: secondary contribution of the European wild apple to the genome of cultivated varieties. PLoS Genetics 8: e1002703.

Coyne, J., and H.A. Orr. 2004. Speciation. Sinauer Associates Inc., Sunderland, MA.

Crane, J.H., and C.W. Campbell. 1994. The Mango. University of Florida Institute of Food and Agricultural Sciences, Gainesville, FL (pp. 1–7).

Dillon, N.L., I.S.E. Bally, C.L. Wright, L. Hucks, D.J. Innes, and R.G. Dietzgen. 2013. Genetic diversity of the Australian national mango genebank. Scientia Horticulturae 150: 213–226.

Doyle, J.J., and J.L. Doyle. 1990. Isolation of plant DNA from fresh tissue. Focus 12: 13–15.

Earl, D.A., and B.M. vonHoldt. 2012. STRUCTURE HARVESTER: A website and program for visualizing STRUCTURE output and implementing the Evanno method. Conservation Genetics Resources 4: 359–361.

Eaton, D.A.R. 2014. PyRAD: Assembly of de novo RADseq loci for phylogenetic analyses. Bioinformatics 30: 1844–1849.

Ellstrand, N.C., R. Whitkus, and L.H. Rieseberg. 1996. Distribution of spontaneous plant hybrids. Proceedings of the National Academy of Sciences of the United States of America 93: 5090–5093.

Evanno, G., S. Regnaut, and J. Goudet. 2005. Detecting the number of clusters of individuals using the software STRUCTURE: A simulation study. Molecular Ecology 14: 2611–2620.

Falush, D., M. Stephens, and J.K. Pritchard. 2003. Inference of population structure using multilocus genotype data: linked loci and correlated allele frequencies. Genetics 164: 1567– 87.

FAO (2003) Tropical Fruits. In Medium-term prospects for agricultural commodities. Food and Agriculture Organization of the United Nations, Rome, Italy. (pp. 1–6).

FAOSTAT (2013). Food and Agriculture Organization of the United Nations Statistics Division. Available from: http://www.fao.org/faostat/.

Gompert, Z., and C.A. Buerkle. 2010. Introgress: a software package for mapping components of isolation in hybrids. Molecular ecology resources 10: 378–84.

Griffith, W. 1854. Dicotyledonous Plants. In J. McLelland [ed.], Notulae ad Plantas Asiaticas, 414–420. Charles A. Serrao, Calcutta.

Harrison, R.G., and E.L. Larson. 2014. Hybridization, introgression, and the nature of species boundaries. Journal of Heredity 105: 795–809.

Hou, D. 1978. Anacardiaceae. In Flora Malesiana Vol 8 Part 3, 395–548. Sijthoff & Noordhoff International Publishers, Alphen Aan Den Rijn, The Netherlands.

Hubisz, M.J., D. Falush, M. Stephens, and J.K. Pritchard. 2009. Inferring weak population structure with the assistance of sample group information. Molecular Ecology Resources 9: 1322–32.

Huson, D.H., and D. Bryant. 2006. Application of phylogenetic networks in evolutionary studies. Molecular biology and evolution 23: 254–67.

IUCN. 2012. World Conservation Monitoring Centre 1998. IUCN Red List of Threatened Species. Version 2012.2. Available at: www.iucnredlist.org.

Iyer, C.P.A., and R.J. Schnell. 2009. Breeding and genetics. In R.E. Litz [ed.], The mango: Botany, production and uses, 68–83. CAB International, Cambridge.

Jakobsson, M., and N.A. Rosenberg. 2007. CLUMPP: A cluster matching and permutation program for dealing with label switching and multimodality in analysis of population structure. Bioinformatics 23: 1801–1806.

Jombart, T. 2008. Adegenet: A R package for the multivariate analysis of genetic markers. Bioinformatics 24: 1403–1405.

Jombart, T., and I. Ahmed. 2011. adegenet 1.3-1: New tools for the analysis of genome-wide SNP data. Bioinformatics 27: 3070–3071.

Kamvar, Z.N., J.F. Tabima, and N.J. Grünwald. 2014. *Poppr*: an R package for genetic analysis of populations with clonal, partially clonal, and/or sexual reproduction. PeerJ 2: e281.

Kiew, R. 2002. Mangifera odorata (Anacardiaceae) is a hybrid. 54: 205–206.

Kostermans, A.J.G.H., and J.M. Bompard. 1993. The mangoes: Their botany, nomenclature, horticulture, and utilization. Academic Press, San Diego, CA.

Kuhn, D.N., I.S.E. Bally, N.L. Dillon, D. Innes, A.M. Groh, J. Rahaman, R. Ophir, et al. 2017. Genetic map of mango: A tool for mango breeding. Frontiers in Plant Science 8: 1–11.

De Langhe, E., E. Hribová, S. Carpentier, J. Dolezel, and R. Swennen. 2010. Did backcrossing contribute to the origin of hybrid edible bananas? Annals of Botany 106: 849–57.

Lim, T.K. 2012a. Mangifera laurina. In Edible medicinal and non-medicinal plants, 124–126. Springer Netherlands, Dordrecht, New York.

Lim, T.K. 2012b. Mangifera odorata. In Edible medicinal and non-medicinal plants, 127–130. Springer Netherlands, Dordrecht, New York.

Mallet, J. 2007. Hybrid speciation. Nature 446: 279–83.

Miller AJ, Gross BL (2011) From forest to field: perennial fruit crop domestication. American Journal of Botany 98(9), 1389–1414.

Mukherjee, S.K. 1949. The mango and its wild relatives. Science and Culture 26: 5–9.

Mukherjee, S.K., and R.E. Litz. 2009. Introduction: Botany and importance. In R. E. Litz [ed.], The mango: Botany, production and uses, 1–18. CAB International, Wallingford, UK.

Peterson, B.K., J.N. Weber, E.H. Kay, H.S. Fisher, and H.E. Hoekstra. 2012. Double digest RADseq: An inexpensive method for de novo SNP discovery and genotyping in model and non model species. PloS one 7: 1–11.

Petit, R.J., and A. Hampe. 2006. Some evolutionary consequences of being a tree. Annual Review of Ecology, Evolution, and Systematics 37: 187–214.

Pritchard, J.K., M. Stephens, and P. Donnelly. 2000. Inference of population structure using multilocus genotype data. Genetics 155: 945–59.

Rieseberg, L.H. 1997. Hybrid origins of plant species. Annual Review of Ecology and Systematics 28: 359–389.

Rodríguez, F., M. Ghislain, A.M. Clausen, S.H. Jansky, and D.M. Spooner. 2010. Hybrid origins of cultivated potatoes. Theoretical and Applied Genetics 121: 1187–98.

Rosenberg, N.A. 2004. Distruct: a program for the graphical display of population structure. Molecular Ecology Notes 4: 137–138.

Schnell, R.J., C.T. Olano, W.E. Quintanilla, and A. W. Meerow. 2005. Isolation and characterization of 15 microsatellite loci from mango (*Mangifera indica* L.) and cross-species amplification in closely related taxa. Molecular Ecology Notes 5: 625–627.

Servedio, M.R., and M. a. F. Noor. 2003. The role of reinforcement in speciation: Theory and data. Annual Review of Ecology, Evolution, and Systematics 34: 339–364.

Soltis, P.S., and D.E. Soltis. 2009. The role of hybridization in plant speciation. Annual Review of Plant Biology 60: 561–88.

Stebbins, G.L. 1950. Variation and Evolution in Plants. Columbia University Press, New York, NY.

Teo, L.L., R. Kiew, O. Set, S.K. Lee, and Y.Y. Gan. 2002. Hybrid status of kuwini, *Mangifera odorata* Griff. (Anacardiaceae) verified by amplified fragment length polymorphism. Molecular Ecology 11: 1465–9.

Wang, X., Y. Xu, S. Zhang, L. Cao, Y. Huang, J. Cheng, G. Wu, et al. 2017. Genomic analyses of primitive, wild and cultivated citrus provide insights into asexual reproduction. Nature Genetics 49: 765–772.

Warschefsky, E. J. Evolution and domestication genomics of the mango genus, Mangifera (Anacardiaceae). Florida International University. https://digitalcommons.fiu.edu/etd/3824

Warschefsky, E., R.V. Penmetsa, D.R. Cook, and E.J.B. von Wettberg. 2014. Back to the wilds: Tapping evolutionary adaptations for resilient crops through systematic hybridization with crop wild relatives. American journal of botany 101: 1791–800.

Warschefsky, E.J., L.L. Klein, M.H. Frank, D.H. Chitwood, J.P. Londo, E.J.B. von Wettberg, and A.J. Miller. 2016. Rootstocks: Diversity, domestication, and impacts on shoot phenotypes. Trends in Plant Science 21: 418–437.

Warschefsky, E. J., and E. J. B. Wettberg. 2019. Population genomic analysis of mango *(Mangifera indica)* suggests a complex history of domestication. New Phytologist 222: 2023–2037.

Wu, G.A., J. Terol, V. Ibanez, A. López-García, E. Pérez-Román, C. Borredá, C. Domingo, et al. 2018. Genomics of the origin and evolution of *Citrus*. Nature.

Yakimowski, S.B., and L.H. Rieseberg. 2014. The role of homoploid hybridization in evolution: A century of studies synthesizing genetics and ecology. American Journal of Botany 101: 1247–1258.

Yamanaka, N., M. Hasran, D.H. Xu, H. Tsunematsu, S. Idris, And T. Ban. 2006. Genetic relationship and diversity of four *Mangifera* species revealed through AFLP analysis. Genetic Resources and Crop Evolution 53: 949–954.

