## Supplemental Figures S1-S4 for "The genetic composition of hybrid *Mangifera*"

### Supplementary Information

**Figure S1.** Plot of $\Delta K$ metric as described by Evanno et al. (2005) for the *M. odorata* dataset. The optimal number of ancestral populations is deemed to be that which has the highest value of $\Delta K$.

**Figure S2.** Plot of expected distribution of rbarD for unlinked loci using 999 permutations (grey bars) and actual rbarD for *M. odorata* samples.

**Figure S3.** Plot of $\Delta K$ metric as described by Evanno et al. (2005) for the *M. casturi* dataset. The optimal number of ancestral populations is deemed to be that which has the highest value of $\Delta K$.

**Figure S4** Plot of expected distribution of rbarD for unlinked loci using 999 permutations (grey bars) and actual rbarD for **(A)** *M. casturi* and **(B)** *M. quadrifida* samples.

**Figure S1.**

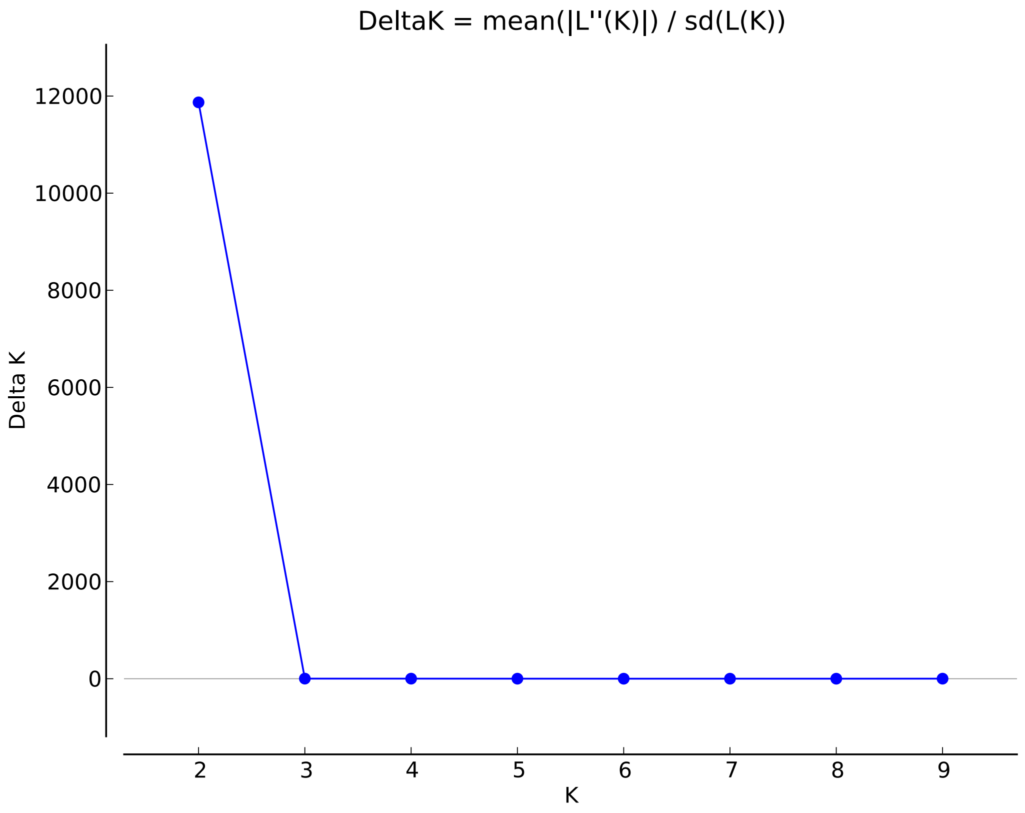
DeltaK = mean(|L''(K)|) / sd(L(K))
Delta K
12000
10000
8000
6000
4000
2000
0
2
3
4
5
6
7
8
9
K

**Figure S2.**

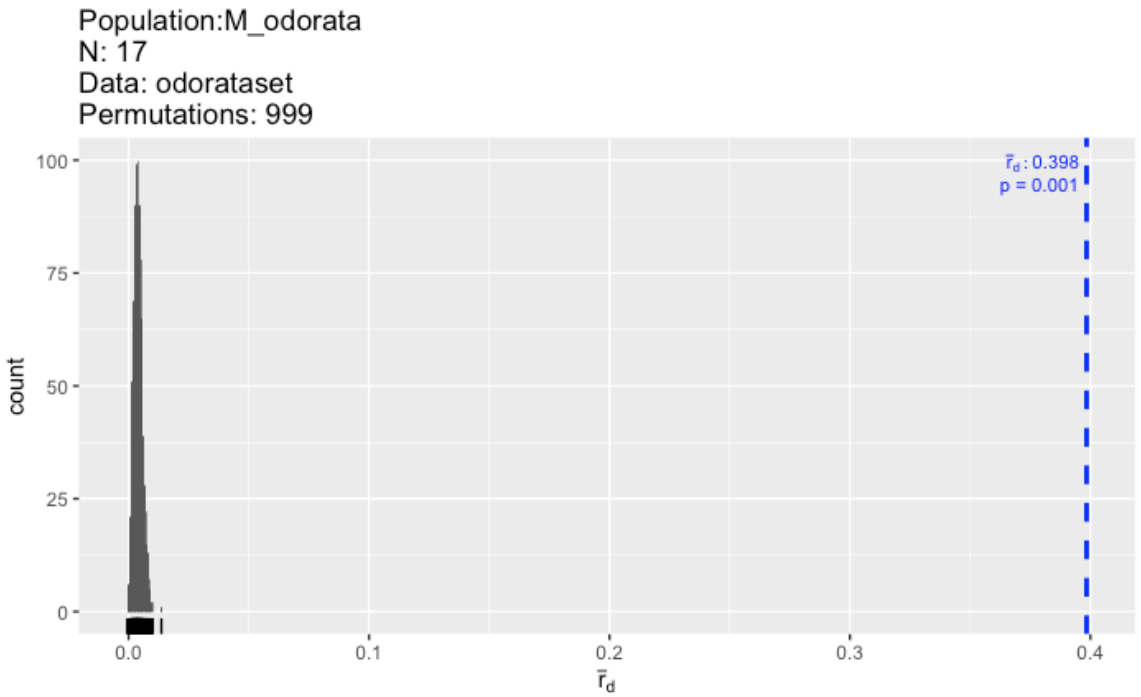
Population:M_odorata
N: 17
Data: odorataset
Permutations: 999
$\bar{r}_d$ : 0.398
p = 0.001
count
100
75
50
25
0
0.0
0.1
0.2
0.3
0.4
$\bar{r}_d$

**Figure S3.**

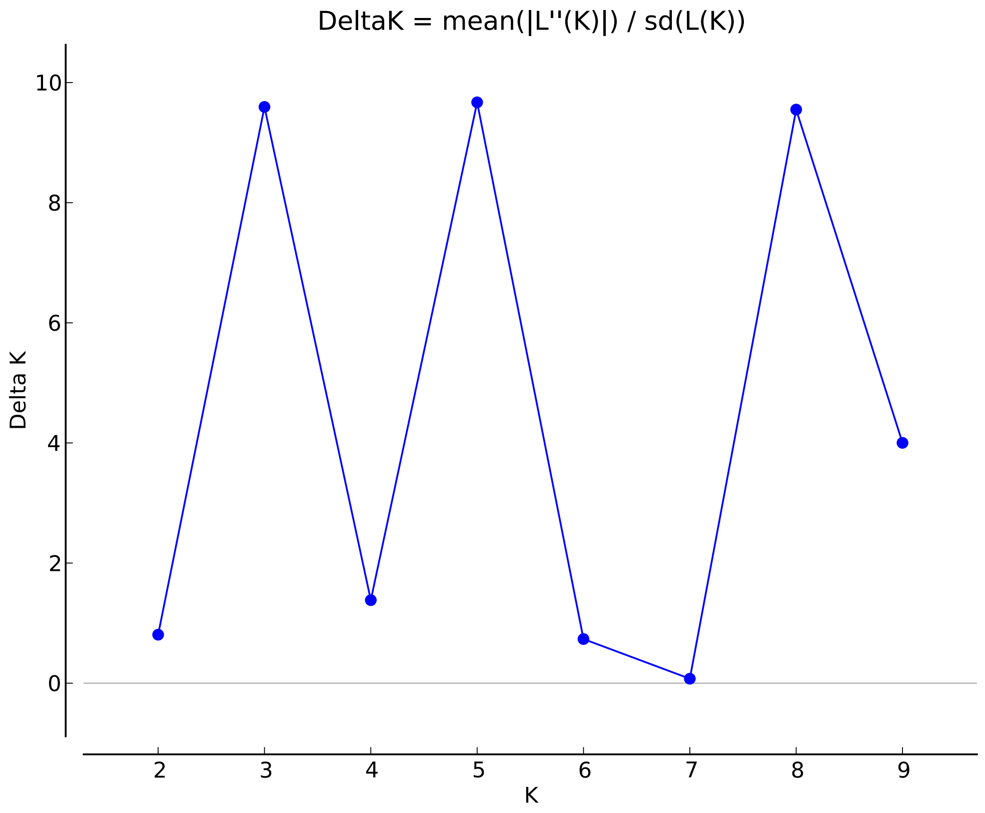

**Figure S4**

**A)**

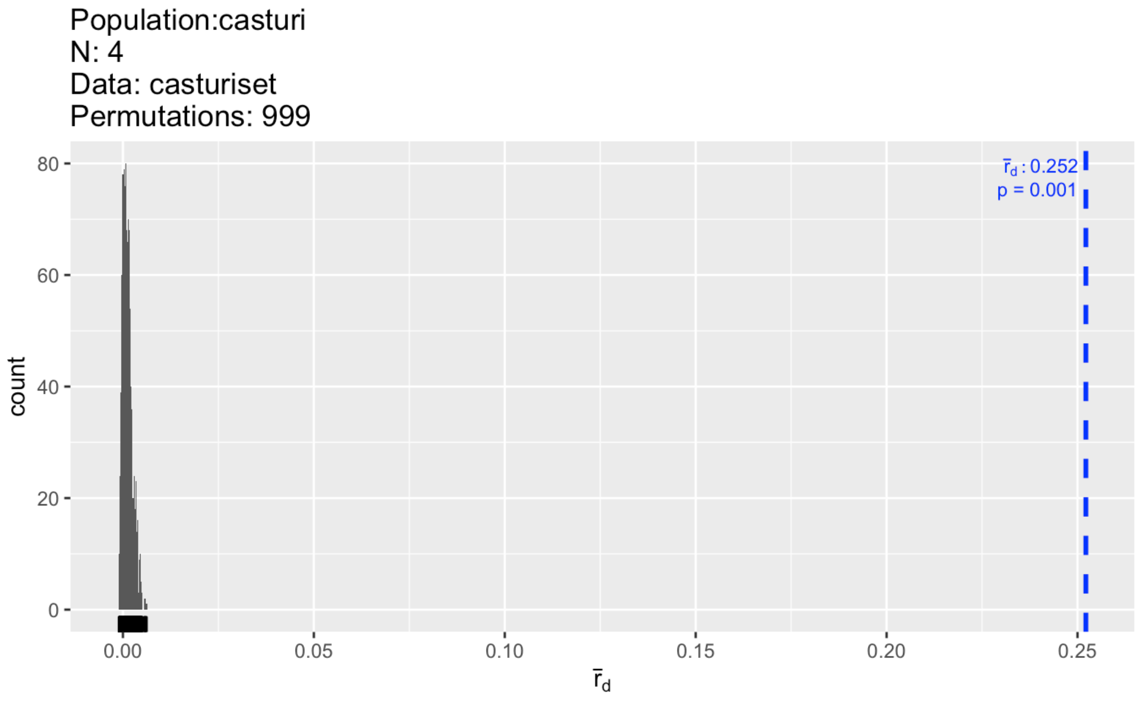
Population:casturi
N: 4
Data: casturiset
Permutations: 999
r̄d : 0.252
p = 0.001
count
80
60
40
20
0
0.00
0.05
0.10
0.15
0.20
0.25
r̄d

**B)**

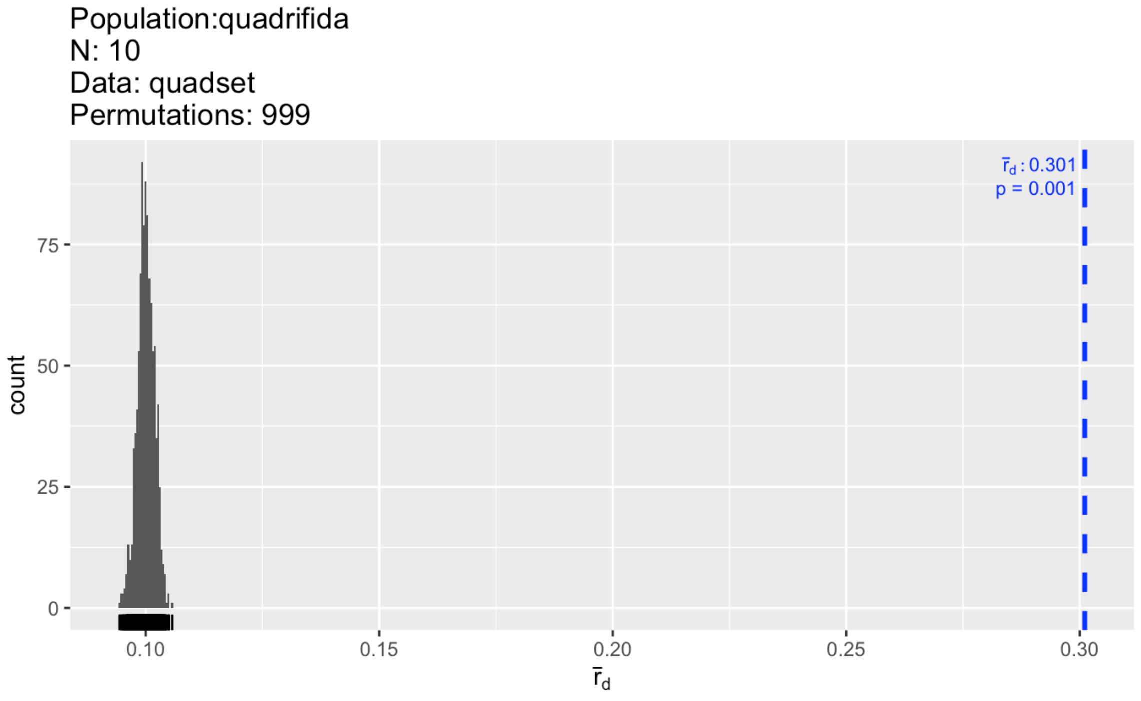
Population:quadrifida
N: 10
Data: quadset
Permutations: 999
r̄d : 0.301
p = 0.001
count
75
50
25
0
0.10
0.15
0.20
0.25
0.30
r̄d
